# Sex-dependent effects of infection on guppy reproductive fitness and offspring parasite resistance

**DOI:** 10.1101/2023.10.29.564622

**Authors:** Isabella L. G. Weiler, Rachael D. Kramp, Faith Rovenolt, Jessica F. Stephenson

**Author notes:** these authors contributed equally to this work (Isabella L. G. Weiler, Rachael D. Kramp).

## Abstract

1. Infection imposes energetic costs on hosts. Hosts typically respond by shifting resources, potentially affecting the quantity and quality of offspring they produce. As the sexes differ in their optimal reproductive strategies, infection of mothers versus fathers may affect offspring quantity and quality in different ways.
2. Here, we test how experimental infection of guppies *Poecilia reticulata* with the ectoparasite *Gyrodactylus turnbulli* affects parental reproductive fitness and offspring parasite resistance. We compared breeding pairs in which one or neither parent had previously been infected.
3. In terms of reproductive fitness, parental infection experience did not affect the size, body condition, or number of offspring produced, but fathers who experienced the heaviest infections produced offspring ∼55 days sooner than average. This result may represent terminal investment by the males most affected by infection, or may indicate that these males have a faster pace of life, investing in reproduction at the expense of parasite defence.
4. We found that offspring age, parental infection experience, and parental infection severity together strongly predicted offspring parasite resistance. Only among pairs in which one parent had been infected, older offspring, which were those born soonest after the parent’s infection, tended to experience heavier infections. This result may therefore reflect temporary infection-induced reductions in parental investment in offspring quality. Beyond this effect of offspring age, offspring of infected mothers experienced 105 fewer worm days than those of infected fathers: fathers, but not mothers, that experienced heavy infections themselves produced offspring that also experienced heavy infections. The parent-offspring regression for infected fathers is therefore consistent with previous evidence that parasite resistance is heritable in this system, and yields a narrow sense heritability estimate of 0.69±0.13. By contrast, the mother-offspring regression (slope: -0.13±0.17) provides novel insight that mothers may engage in transgenerational immune priming.
5. Overall, our results suggest that the sexes strike a different balance between offspring quantity and quality when faced with infection, with potentially broad implications for disease and host-parasite coevolutionary dynamics in nature.

## Introduction

Infected parents may change the quantity and quality of the offspring they produce, both because of the energetic constraints (Tizard, 2008) or pathology (Armour et al., 2020; Billi et al., 2019; Khan et al., 2017) associated with infection, and because the optimal investment in defence (Boots & Bowers, 2004; Graham, 2013; Houston et al., 2007; Tschirren & Richner, 2006; Viney et al., 2005) and reproductive strategy may differ under the threat of infection (Schmid-Hempel, 2003; Sheldon & Verhulst, 1996). Intuitively, parental infection could induce the production of fewer, lower quality offspring due to decreased parental resources (Hagmayer et al., 2020; Vantaux et al., 2014). However, infection can also induce the production of more offspring as infected parents ‘terminally invest’: hosts increase reproductive output at the expense of their own defence against infection in an effort to produce offspring before their own death (Duffield et al., 2017). Infection may also induce the production of higher quality offspring, particularly if they are likely to be exposed to the same parasite, through parental investment in the immunological defences of their offspring, known as “transgenerational immune priming”, or TGIP (Roth et al., 2018). TGIP can last from shortly after birth, (in the case of e.g. maternal antibodies (Breuil et al., 1997; Brown et al., 1997)) to subsequent generations (in the case of epigenetic changes (Beemelmanns, 2019)), depending on the host species and the disease to which they are responding (Roth et al., 2018). TGIP is costly to both parents and offspring, however: manufacturing parentally donated factors such as proteins or antibodies requires resources and energy, and any form of priming may require offspring to mount an immune response, even in the absence of the disease (Roth et al., 2018).

Mothers and fathers typically take different strategies to maximise fitness, even in the absence of infection. At the ultimate level, parental investment theory (Trivers, 1972) posits that as females (in species with typical sex roles) invest more in individual gametes or zygotes, they are limited in the rate at which they can produce offspring. Females therefore maximise fitness by investing in the probability of survival and reproduction of the offspring they produce. Males, by contrast, produce many more gametes at a much higher rate and, all else being equal, maximise fitness by mating with as many females as possible. Therefore, in the absence of infection, males should optimally invest in offspring quantity, and females, quality.

These life history differences between the sexes have implications for their investment in their own immunity (Fischer et al., 2011; Rolff, 2002; Stoehr & Kokko, 2006), and potentially how they shift their parental investment post-infection. Females typically invest more in their immune and self-repair systems to maximise longevity, and thus lifetime reproductive success, whereas males do not experience this same selection on longevity (Rolff, 2002; Stoehr & Kokko, 2006). As well as ultimate drivers, the sexes may differ in the proximate mechanisms through which they can affect offspring quantity or quality. For example, in many systems, mothers have more opportunity to affect the phenotypes of their offspring (Wassarman et al., 2001; Wolf & Wade, 2009), and as a result offspring phenotypes are often strongly correlated with maternal phenotypes, for example in live-bearing fish (Hagmayer et al., 2018) and humans (Zhang et al., 2018). This may hold true for TGIP: although the underlying mechanisms have not been fully elucidated, promising candidates include parentally donated proteins (Adrian-Kalchhauser et al., 2020) that are likely too large to be transferred by sperm (Arnqvist & Rowe, 2005; Wassarman et al., 2001). Indeed, no relationships among antibody titres were detected in vertebrates when fathers but not mothers were primed (Gasparini et al., 2002; Reid et al., 2006). However, other mechanisms, such as epigenetic modifications (i.e., altered immune gene expression in offspring (Vilcinskas, 2021)) or nonnuclear effects (Simms & Triplett, 1996), could be mediated by either parent. Indeed, a recent meta-analysis of TGIP in invertebrates found that the sex of the parent primed did not affect the strength of TGIP (Rutkowski et al., 2023). Such mechanisms likely function among vertebrates, too: among pipefish, both offspring and grand-offspring of primed parents show altered gene expression (Beemelmanns & Roth, 2017).

However, more broadly, work in vertebrates has historically been heavily biased towards investigating maternal effects (Roth et al., 2012, 2018). Indeed, evidence for paternally-mediated TGIP in vertebrates is limited to sex-role reversed species (seahorses and pipefish biparental TGIP influences offspring lymphocyte proliferation and immune gene expression (Beemelmanns & Roth, 2016, 2017; Roth et al., 2012)), or those with some paternal care (Pizzolon et al., 2010).

Here, we test how infection with *Gyrodactylus turnbulli* affects the reproductive fitness of male and female guppies, and the quality of the offspring produced. This highly tractable and manipulable system has been used extensively (Harris & Lyles, 1992; Madhavi & Anderson, 1985; Stephenson et al., 2018; Walsman et al., 2022), and has characteristics that make it particularly appropriate for understanding transgenerational effects. Indeed, guppy exposure to predators has transgenerational effects mediated by both parents (Stein & Hoke, 2022). For infection, resistance to *G. turnbulli* is heritable (Dargent et al., 2016; Madhavi & Anderson, 1985), enabling us to partially control for host genotype when investigating other potential sources of resistance. The parasite is associated with lower survival (Stephenson et al., 2016; van Oosterhout et al., 2007) and body condition (Stephenson, van Oosterhout, et al., 2015) of wild guppies, indicating it depletes host resources. Furthermore, guppy immune gene expression is affected by infection with *Gyrodactylus* spp. (Konczal et al., 2020). The key conditions for TGIP are present in this system (Roth et al., 2018): *G. turnbulli* can be directly transmitted from mother to offspring during birth, so offspring of infected mothers are likely to be exposed to the same infection; and *G. turnbulli* primarily reproduce asexually (Schelkle et al., 2012), indicating infections in natural populations have some genotypic stability.

The biology of our system also suggests that the sexes may differ in how infection with *G. turnbulli* affects parental investment. First, the parasite interacts with the host sexes differently: males are less likely to be found infected in nature (Stephenson, Van Oosterhout, et al., 2015), and experience lower infection loads both in the wild (Stephenson, Van Oosterhout, et al., 2015; Walsman et al., 2022) and during experimental infections (Dargent et al., 2016) than females.

Despite this, females may be more likely than males to survive their infections in nature (Clark et al., 2023; Stephenson, van Oosterhout, et al., 2015; van Oosterhout et al., 2007), and appear better able to maintain the same behavioural phenotypes as uninfected fish (Jog et al., 2022; Stephenson, 2019). Female guppies may be well-positioned to prime their offspring: they give birth to live offspring whose eggs are provisioned only at the time of fertilization, and female phenotype affects offspring phenotype (Shikano & Taniguchi, 2005). However, in other fish, maternally-provisioned antibodies often disappear after offspring pass certain developmental stages such as absorbing the yolk sac (Breuil et al., 1997; Brown et al., 1997) which, in livebearers such as the guppy, usually happens before they are born (Norazmi-Lokman et al., 2016). Male guppies may also be able to directly affect offspring phenotype through sperm quality: in an artificial insemination experiment, sperm from males fed a high quality diet was of a higher quality itself, and produced offspring that were heavier at birth (Evans et al., 2017).

Sperm is the only potential route for paternally-derived TGIP in this system: infection may affect a father’s investment in his sperm, which in turn could affect the quality of his offspring, as potentially reflected in their parasite resistance. We also know that infection with *G. turnbulli* affects DNA methylation in guppies (Hu et al., 2018a), raising the possibility of epigenetically-mediated (and thus potentially bi-parental) TGIP. Such epigenetic changes can last generations after an initial immune challenge (Beemelmanns, 2019).

We address two main questions: 1) Does *G. turnbulli* infection affect guppy reproductive fitness? 2) Does parent infection affect offspring quality and response to *G. turnbulli* infection? Based on parental investment theory, we predicted that males should prioritize offspring number, whereas females should prioritise offspring quality, after exposure to a parasite. We quantified offspring number as the total number of offspring a pair produced during our observation period, and latency to birth: together these predict how many offspring a female (and her mate in monogamous pairs, such as in our experiment) may produce in her lifetime, assuming all else is constant. We quantified offspring quality as their size, body condition, and how heavily infected they became during experimental infection.

## Materials and Methods

### (a) Fish Origin and Maintenance

We used laboratory-bred descendants of guppies from the Caura River in Trinidad. This laboratory population was founded by fish collected in 2012 from a high-predation population located at an undeveloped, rural site on the Caura River, Trinidad (UTM: 20 P 679527.7 m E, 1180376.4 m N based on WGS84 Datum; elevation 112m). Since 2012, the population has been maintained in large numbers, and we continue to see substantial variation in disease resistance, as is clear from our results. Fish were maintained and bred in 20 L tanks of mixed sexes, under standard conditions, as follows. Tanks were on a recirculating system (Aquaneering) with approximately 20% daily water changes, a lighting schedule of 12L:12D, and an average temperature of 25±1°C. In these systems, wastewater from tanks passed through foam, sand, and ultraviolet filters before re-entering other tanks, preventing unintentional parasite transmission. Fish were fed flake and Artemia daily.

### (b) Parasite origin and maintenance

Our parasite line was established by transferring one worm from a commercially obtained guppy to an uninfected host from a mixed laboratory-bred stock, a ‘culture fish’. This parasite line was maintained on groups of three to six culture fish per 1.8 L tank under standard conditions. Twice per week, all culture fish were anesthetized (tricaine methanesulfonate ‘MS222’; 4g per L) and parasites were counted using a dissecting microscope. Uninfected fish were added to replace recovered or dead fish.

### (c) Parental infection and breeding design

Parental fish were anaesthetized (using MS222 as above) and placed in close proximity to a dead, highly infected culture fish until at least two worms transmitted (mean±SEM=2.8±0.27), as observed under a dissecting microscope. Once infected, parental fish were isolated in 1.8 L tanks maintained under standard conditions for the duration of their infection. We counted worms every three days for 15 days to record individual infection trajectory over time. After 15 days, we treated fish with levamisole (0.002g per L) to clear infection. We verified that this treatment had cleared infection by screening the fish for parasites, under anaesthetic, three times with a minimum of four days between screens. We recorded the length and weight of the parents before and after infection. Parasite naïve parents were exposed to the same conditions and manipulations, with the exception of parasites.

We set up 38 total breeding pairs of one male and one female using the recovered fish and parasite naïve fish in three different treatments: recovered mother, naïve father (13 pairs); recovered father, naïve mother (11 pairs); and both parents naïve (14 pairs). We did not set up pairs in which both parents were recovered because previous experience suggested this treatment may not have yielded sufficient offspring, and our primary objective was to examine the independent effects of maternal and paternal infection. These breeding pairs were housed in 1.8 L tanks under standard conditions, including fry mesh structures and gravel to reduce possible fry cannibalism by the parents. Between May and December 2021, these breeding tanks were checked daily for the presence of offspring, which were moved to a separate 1.8 L tank. Each brood produced from a pair was moved to a separate 1.8 L tank.

### (d) Offspring infection

Once they reached adulthood, offspring were separated into individual 1.8 L tanks. A total of 108 offspring were collected: 32 from recovered fathers and naïve mothers; 38 from recovered mothers and naïve fathers; and 38 from naïve parents. We used two batches of fish, one in August 2021 (52 offspring from 33 breeding pairs, with an average age at infection of approximately 4.4 months) and one in May 2022 (56 offspring from 32 breeding pairs, with an average age at infection of approximately 7.8 months). According to the needs of other concurrent experiments, 94 fish were infected manually, as described under parental infection, and 14 were infected via “co-housing”: they were housed in 1.8 L tanks with an infected conspecific and screened for infection every 24 hours. Once infected, these fish were isolated in a 1.8 L tank and treated the same as the manually infected offspring and parents from that point forwards. As noted below, we included infection type as a fixed effect in the statistical models and found that it did not explain significant variation in any of our response variables. On all infected offspring, we counted the worms every three days for 12 days and then sacrificed the host.

### (e) Data Analysis

We used R statistical software (R Core Team, 2020) to analyse the data, and provide the code and output in the supplement. We used the data from this experiment to address two main questions: 1) Does infection affect reproductive fitness? 2) Does parent infection affect offspring response to infection? We addressed each question using a) data from all pairs, whether or not one parent had been infected; and b) only data from pairs in which one parent had been infected. This enabled us to evaluate the extent to which parental exposure to infection affected reproductive fitness and offspring infection (a); and the extent to which the intensity of the parent’s infection explained variation in these two outcomes (b). As our measure of the parent’s infection intensity, and offspring response to infection, we used the area under the curve of infection load over time (‘infection integral’).

As our metrics of reproductive fitness, we used: the latency between the parent’s infection and the birth of the offspring; the total number of offspring a pair produced during the observation period; and the body length and body condition of those offspring at adulthood. We used each of these four metrics as the response variable in separate linear (mixed) models (Gaussian error family, identity link function) in glmmTMB (Brooks et al., 2017) using first the data from all pairs, and then data from only pairs in which one parent had been infected (i.e., a total of eight models). Each of these eight models included as fixed effects the parental infection treatment (mother, father, or neither parent infected), mother’s body condition, mother’s body length, and father’s body condition. For response variables relevant to the level of the individual offspring (birth latency, body length, body condition), the models included mother identity as a random factor as many pairs produced multiple offspring. Because male and female guppies typically differ in body condition metrics and length, we included sex of the offspring as a fixed effect in those models. For the total number of offspring models, we used only one observation per pair. The four models using only data from pairs in which one parent had been infected additionally included the (log transformed) infected parent’s infection integral and the interaction with parental infection treatment as fixed effects.

We used two separate linear mixed models to test how a) parental exposure to infection, and b) the severity of that infection, affects offspring response to infection (Gaussian error family, identity link function in glmmTMB (Brooks et al., 2017)). Both models included as fixed effects the sex, body condition, body length (residuals on sex), age at infection, the method of experimental infection (manual vs. cohoused), and the number of parasites establishing an infection on the offspring. Both models also included the parental infection treatment (mother, father, neither), and the interaction between parental infection treatment and offspring age at infection. As random effects, both models included the mother’s identity to control for the non-independence of data from multiple offspring from the same pair. In addition, the model including parental infection severity included parental infection integral (log transformed), and its interaction with parental infection treatment, plus a three-way interaction with these two variables and offspring sex. We checked the fit of all models by visually assessing the plots and tests produced by the DHARMa package ((Hartig, 2022) provided in the supplement). DHARMa simulates standardized residuals to thoroughly validate complex mixed models such as those we use here. The figures were plotted using visreg (Breheny & Burchett, 2017) and ggplot2 (Wickham, 2016).

We estimated narrow-sense heritability from the slope of the parent-offspring regression (Åkesson et al., 2007, 2008; Falconer & Mckay, 1997). We rescaled parent and offspring infection integrals and included offspring age at infection (which in our main analysis explained a highly significant portion of the variation in offspring infection integral) as a fixed effect in our models, and mother ID as a random effect to account for the fact that pairs contributed more than one offspring. We used glmmTMB (Brooks et al., 2017) to estimate slope, standard error, and probability the slope differed from 0.

## Results

### We did not find concerning issues with the fits of our models and present the plots and tests of assumptions in the supplement

#### (a) Does infection affect reproductive fitness?

We found that previously infected and parasite naïve guppies produced the same number of offspring, at the same rate, and these did not differ in either size or body condition at adulthood (all *P* > 0.25, see supplement).

Similarly, we found that the intensity of parental infection did not significantly affect the number of offspring a pair produced, or the size or body condition of those offspring at adulthood (*P* = 0.06-0.97, see supplement). One of these results was marginally non-significant: parents that experienced heavier infection loads produced offspring with higher body condition at adulthood (Chisq = 3.51, df = 1, *P* = 0.061).

We found that the number of days between parental infection and offspring birth depended on the intensity of that infection, and this relationship differed between infected mothers and fathers (Fig. 1; interaction between parental infection treatment and integral: Chisq = 5.13, df = 1, *P* = 0.024). We reran this analysis on the data split by sex and found that the correlation among fathers differed significantly from 0 (effect of infection integral: Chisq = 5.49, df = 1, *P* = 0.019), whereas that among mothers did not (Chisq = 0.87, df = 1, *P* = 0.35).

**Fig. 1:**
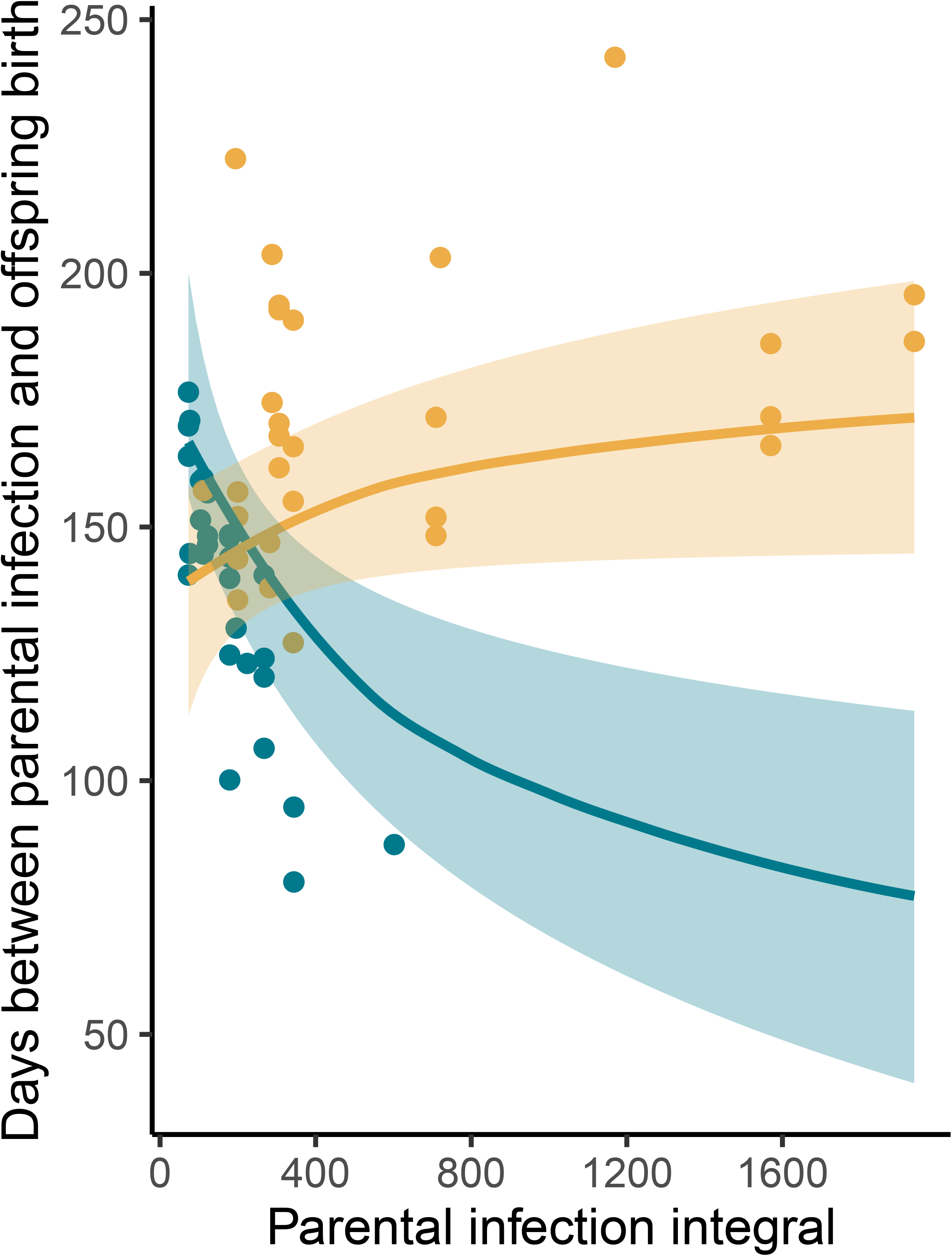
Pairs in which fathers (darker blue) had more intense infections produced broods quicker, whereas the intensity of the mother’s (lighter yellow) infection did not affect birth latency. Points represent the partial residuals from the model described in the main text, lines give the model fits, and shading the standard error. We quantified infection intensity using ‘infection integral’, or the area under the curve of infection load over time.

#### (b) Does parent infection affect offspring response to infection?

We found that the effect of parental infection treatment on offspring response to infection depended on the age of the offspring at infection, and the parental infection treatment: among offspring with a previously infected parent, younger offspring experienced lower infection intensities than older offspring (Fig. 2; interaction between parental infection treatment and offspring age at infection: Chisq = 7.39, df = 2, *P* = 0.025). We also found that the more parasites initially established on the fish, the higher their infection integral (Chisq = 9.69, df = 1, *P* = 0.002). No other variables, including offspring sex, explained any of the variation in offspring infection integral (see supplement for full model output). Our *post-hoc* tests of the correlation in Fig. 2 show that it is significantly different from 0 for offspring of infected fathers (Chisq = 16.57, df = 1, P < 0.000) and mothers (Chisq = 4.70, df = 1, *P* = 0.03), but not those of parasite naïve parents (Chisq = 2.39, df = 1, *P* = 0.122; see supplement).

**Fig. 2:**
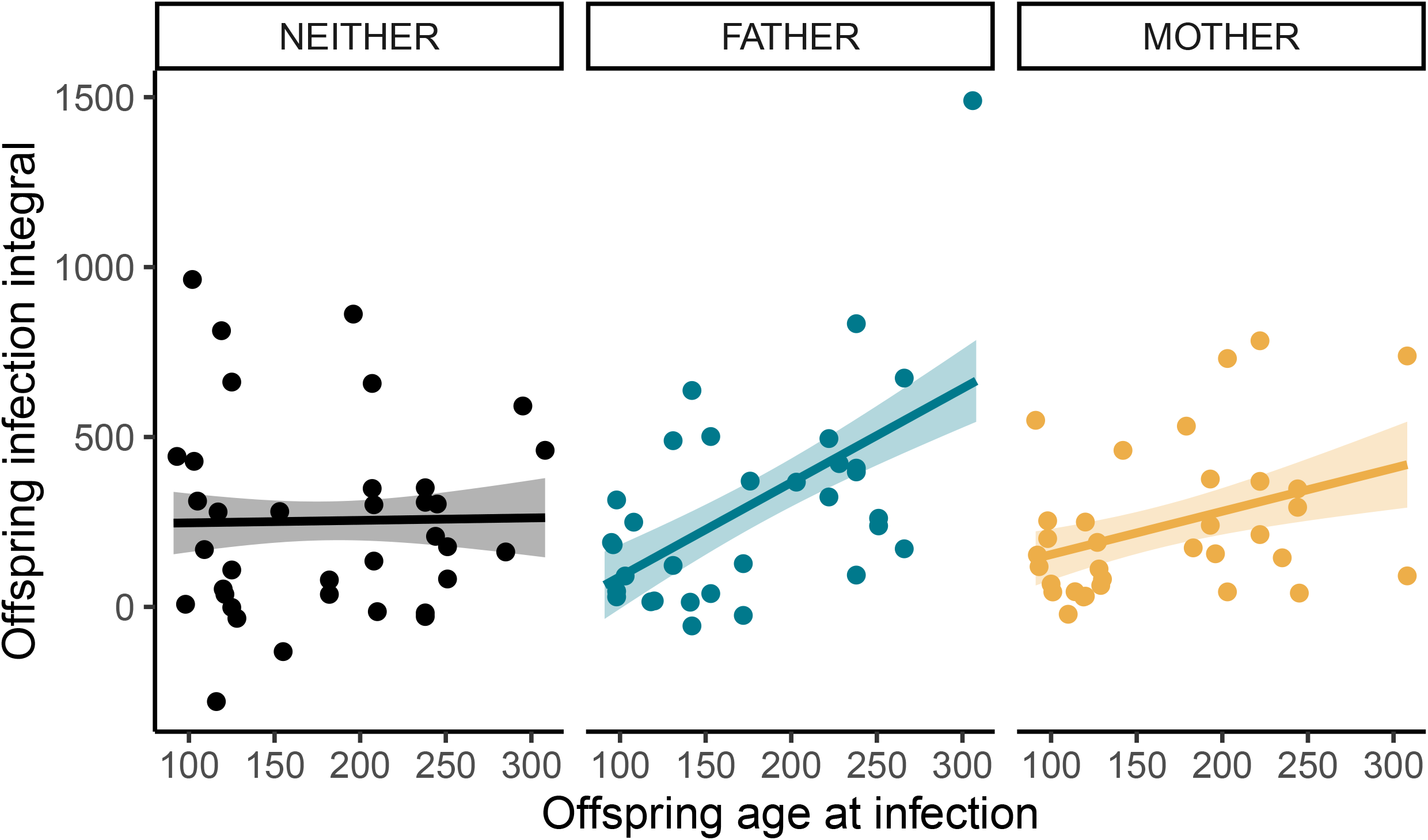
Among offspring with an infected parent, older fish tended to experience more intense infections, but no such relationship exists among offspring of two parasite-naïve parents. Points represent the partial residuals from the model described in the main text, lines give the model fits, and shading the standard error. We quantified infection intensity using ‘infection integral’, or the area under the curve of infection load over time.

Among offspring with one infected parent, we found their infection integral was significantly predicted by whether the mother or the father had been infected, but this effect depended both on offspring age at infection, and the intensity of the parent’s infection. As in the model including all offspring, younger offspring experienced lower infection intensities than older offspring (Chisq = 19.74, df = 1, *P* < 0.000). We also found a significant correlation between the father’s infection integral and that of the offspring, but no significant correlation between mother and offspring infection integral (Fig 4.; interaction between parental infection integral and parental infection treatment: Chisq = 11.89, df = 1, *P* = 0.0006). No other variable explained a significant portion of the variation (see supplement), including the interaction between parental infection treatment and offspring age: the correlations in Fig. 2 do not therefore differ significantly between offspring with infected mothers and fathers. Our *post-hoc* tests confirmed that the correlation between father and offspring infection integrals differs from 0 (Chisq = 13.07, df = 1, *P* = 0.0003), whereas that between mothers and offspring does not (Chisq = 0.63, df = 1, *P* = 0.43).

#### (c) What is the heritability of parasite resistance?

The slope of the father-offspring regression is 0.69±0.13 (estimate±standard error; *P* < 0.00001), and 0.53±0.18 (*P* = 0.003) after the removal of one outlier. The slope of the mother-offspring regression is -0.13±0.17 (*P* = 0.45).

## Discussion

Our results show that parental infection does affect the quantity and quality of offspring a pair produces, and that these effects depend on which parent was infected. We found that males that experienced the heaviest infections produced offspring ∼55 days quicker than average, which may result in these males producing more offspring over their lifetimes. We also found that older offspring experienced heavier infections, but only among those produced by parents with experience of infection. Finally, we found that offspring of fathers who experienced heavy infections also experienced heavy infections, whereas offspring of mothers showed no such pattern. On average, offspring of infected mothers experienced 105 fewer worm days than offspring of infected fathers. Overall, our results indicate how parasites may change reproductive investment strategies, and how this affects the next generation of hosts. Here we discuss each of our results in turn.

Our observation that fathers that become heavily infected produced offspring more quickly (Fig. 1) is consistent with other observations from this system. First, these males may mate more quickly than those that experienced lighter infections and thus their broods are produced earlier. Previous work has shown that *Gyrodactylus* spp. infection (Kennedy et al., 1987; López, 1998, 1999) and immune challenges (Encel et al., 2023) affect guppy reproductive behaviour and, consistent with this explanation, the heaviest infected males appear more interested in contact with females (Stephenson, 2019). This behavioural explanation may be a component of a broader explanation that fathers who become heavily infected may generally have a faster pace of life. Such males would bias investment towards reproduction, including reproductive behaviours, and away from immune investment, resulting in their heavier infection loads. There is evidence from natural guppy populations that pace of life does predict resistance to *Gyrodactylus* spp. (Stephenson, van Oosterhout, et al., 2015). Alternatively, rather than being indicative of a pace of life syndrome, this result could indicate that the males that experienced the heaviest infections are terminally investing in reproduction (Duffield et al., 2017): while suggestive results have been reported (López, 1998; Stephenson, 2019) this phenomenon remains to be conclusively demonstrated in guppies.

We found that older offspring experienced heavier infections, but only among those produced by parents with experience of infection (Fig. 2). It is well known that many immune components wane with age across animal taxa (Metcalf et al., 2020; Peters et al., 2019), but the offspring in our experiment were young adults, and the effect of age was absent from those born to parasite naïve parents. We therefore suggest that this result is evidence that parental infection affects investment in offspring quality. Indeed, investment in offspring can respond to the environment among guppy mothers (Cunningham & Russell, 2000; Reynolds & Gross, 1992) and fathers (Evans et al., 2017). The oldest offspring in our experiment are those produced the soonest after parental infection, so could therefore represent those that received least resources from parents depleted by that infection. However, the mechanism underlying this potential explanation is unclear. As maternal antibodies, mRNAs, and proteins are typically absent from guppy offspring within a few days post-birth (Breuil et al., 1997; Brown et al., 1997; Norazmi-Lokman et al., 2016), and such proteins are too large to be transferred by sperm (Arnqvist & Rowe, 2005; Wassarman et al., 2001), it seems unlikely this result is due to waning TGIP.

We found that offspring infection intensity was strongly correlated with parental infection intensity among offspring of infected fathers, but not among those of infected mothers, even after controlling for the effect of offspring age (Fig. 3). Among offspring of infected fathers, the positive parent-offspring regression we observe is consistent with previous findings that resistance to *Gyrodactylus* spp. is heritable in guppies (Dargent et al., 2016; Madhavi & Anderson, 1985; Phillips et al., 2018), and other fish (Gilbey et al., 2006). Our estimates of narrow-sense heritability from these regressions (0.69 or 0.53 after outlier removal) are high, but comparable to heritability estimates for male guppy ornaments (Morris et al., 2020; van der Bijl et al., 2023), which are correlated with *G. turnbulli* resistance in our focal population (Stephenson et al., 2020).

**Fig. 3:**
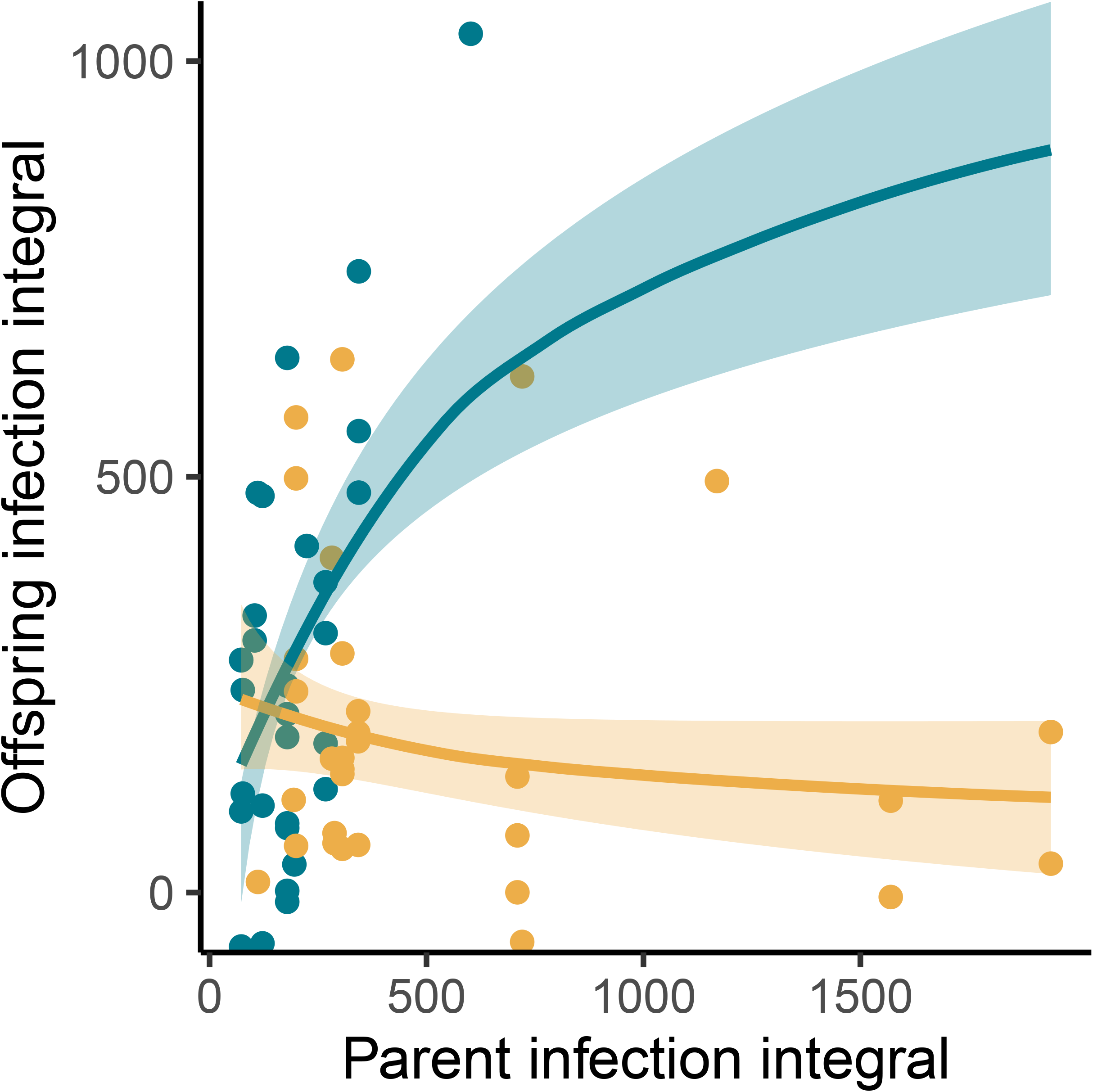
A father’s infection intensity significantly predicts the infection intensity that his offspring experience, but no such relationship exists between the infection intensity experienced by a mother and that of her offspring. Points represent the partial residuals from the model described in the main text, lines give the model fits, and shading the standard error. We quantified infection intensity using ‘infection integral’, or the area under the curve of infection load over time.

Despite the heritability of resistance to *Gyrodactylus* spp., we did not observe a significant mother-offspring regression. Potential mechanisms as to why offspring resembled their fathers but not their mothers in this trait include sex-linked inheritance, autosomal dominant traits, and maternally derived TGIP. We believe maternally derived TGIP provides the most parsimonious explanation. For example, if resistance to *Gyrodactylus* spp. is Y-linked, and thus passed from fathers to sons, the sex of the offspring from infected fathers should explain variation in offspring parasite resistance but it does not. However, there is some evidence that parasite resistance evolves along sex-specific trajectories in natural conditions (Dargent et al., 2016). Autosomal dominant traits also seem an unlikely explanation as males were randomly assigned across our 38 pairs. Furthermore, the Major Histocompatibility Complex (MHC) is important in guppy resistance to these parasites (Fraser et al., 2010; Fraser & Neff, 2009; Phillips et al., 2018), and many MHC genes are codominantly expressed (Herdegen-Radwan et al., 2020). We therefore suggest that maternally derived TGIP most likely underlies our result, though it is clear more work is needed to fully elucidate the mechanisms underlying the difference we observed between father-offspring and mother-offspring regressions.

Our data are therefore consistent with maternally mediated TGIP and provide some insight into potential mechanisms. Offspring of mothers that experienced severe infections had light infections, comparable to those of offspring of mothers who experienced less severe infections. Thus, any parasite load may be sufficient to induce TGIP to the same protective level, or perhaps TGIP is compensatory to load and thus allows all offspring of infected mothers to benefit from a similar level of resistance. The offspring were all adults, several months old, before their infections, which could rule out mechanisms observed in other systems such as endocrine signalling (Kvist et al., 2020) or protein transfer (Adrian-Kalchhauser et al., 2020) from the mother to the offspring. Epigenetic mechanisms would persist in the offspring until adulthood and beyond, and maternal stress can induce epigenetic changes, such as histone modifications and DNA methylation, during gametogenesis in mammals (Gapp et al., 2014).

Indeed, epigenetic changes in response to infection have been observed in female guppies (Hu et al., 2018b) and these changes may be conserved from generation to generation (Heard & Martienssen, 2014). These changes persist after fertilization, could permanently alter offspring immune function, and provide an evolutionary advantage as females can prepare their offspring for the parasites they are likely to encounter in their environment (Willis et al., 2021). Epigenetic changes could also lead to non-immune changes like increased epidermal thickness, which has been implicated in guppy response to stress and interactions with *G. turnbulli* (Gheorghiu et al., 2006, 2009, 2012).

Our results suggest more complexity in the interactions between host and parasite in natural communities. In our system, Trinidadian guppy populations differ in their prevalence of *Gyrodactylus* spp. and the mean intensity of infection, and these are likely not stable through time (Clark et al., 2023; Stephenson, Van Oosterhout, et al., 2015; Walsman et al., 2022). In populations and at times when prevalence is high, our findings suggest that more mothers may prime their offspring, who would develop overall lower infection loads, potentially reducing transmission (Stephenson et al., 2017), and thus affecting population-level prevalence. If maternally derived TGIP has a measurable impact on disease dynamics in nature, it may therefore slow the evolution of host resistance by reducing the strength of parasite-mediated selection. However, host defences do appear to evolve rapidly in response to these parasites (Dargent et al., 2016; Phillips et al., 2018). Heavily infected guppies may not successfully breed in nature, which would erode the effects of TGIP on resistance evolution. Indeed, observed natural infection loads tend to be low (Stephenson, Van Oosterhout, et al., 2015), and patterns of aggregation show that heavily infected hosts are missing from our surveys, presumed dead (Clark et al., 2023). Similarly, the impact of TGIP may be overwhelmed by the fact that males are able to sire offspring (that inherit their parasite resistance) up to 10 months posthumously (Lopez-Sepulcre et al., 2013), and engage in coercive mating so will likely mate (Magurran, 2005), even though females discriminate against males who are infected (Kennedy et al., 1987), or those who may lack resistance (Stephenson et al., 2020). How the TGIP we observe here affects dynamics in nature remains unclear, therefore, but the magnitude of the effect we observe should mean it warrants future investigation.

More broadly, our comparison of the maternally and paternally derived transgenerational effects of infection contributes novel insight valuable across taxa. In accordance with parental investment theory, we found that males appear to prioritise offspring quantity, and females quality, in the face of infection. We also show that maternal and paternal contributions to offspring immunity can differ significantly. Thus, our observed transgenerational effects of infection potentially influence disease dynamics and host-parasite co-evolutionary dynamics in natural communities (McIntire et al., 2023).

## Supporting information

Complete Supplement 1

## Acknowledgements

We thank D. Clark, L. Colgan, D. Henry, M. Janecka, R. Patrick, J. Walsman, C. Wynne for technical assistance. This work was approved by the University of Pittsburgh’s Institutional Animal Care and Use Committee (IACUC), protocol 21069471, and supported by the University of Pittsburgh, NSF DGE: 1747452 (to FR), and an HHMI Gilliam Fellowship (to RDK).

## Data availability

The data are available in the supplement and, upon manuscript acceptance, will be deposited in Dryad and the doi included here. The code and output we used to analyse the data is included in the supplement.

## CRediT authorship statement

Conceptualization = JFS; Data curation = ILGW, RDK, FR; Formal analysis = JFS; Funding acquisition = FR, JFS; Methodology = ILGW, RDK, FR; Project administration = JFS; Resources = JFS; Visualization = JFS; Writing – original draft = ILGW, JFS; Writing – review & editing = ILGW, RDK, FR, JFS. All authors approved gave approval for publication and agreed to be accountable.

