## Supplementary material for "Sex-dependent effects of infection on guppy reproductive fitness and offspring parasite resistance": Complete Supplement 1

#### Introduction

In this paper we test how the fitness of laboratory-bred guppies *Poecilia reticulata* is affected by infection with *Gyrodactylus turnbulli*. Parent fish were experimentally infected, and we counted worms every three days for 15 days. After 15 days, we treated fish with levamisole to clear infection. Recovered fish were paired with naive (uninfected) fish to produce a breeding pair of one male and one female. Naive fish were also paired with other naive fish as a control treatment. Pairs were housed in 1.5 L tanks such that the offspring collected had known parentage. Offspring were collected and housed individually, and then experimentally infected following the same protocol as the infection on their parents. In our analysis, we used data provided in the file `TGIPData.csv`, which contains the following variables about the infected offspring:

**SEX:** Sex of the fish
**INFTYPE:** Whether the fish was infected under anaesthesia through the manual exposure to a heavily infected, euthanized donor fish ('manual'), or through cohousing with a live infected donor ('cohouse')
**PREL:** Fish length in mm before infection
**LENGTHRESID:** The residuals of length of the focal fish on sex. Females tend to be larger than males: using the residuals of this relationship allows us to test for both size and sex differences.
**PREW:** Fish weight in mg before infection
**LASTW:** Fish weight in mg after infection
**preSMI:** Body condition - scaled mass index - before infection

**postSMI:** Body condition - scaled mass index - after infection
**deltaSMI:** Body condition - scaled mass index - change over the course of infection
**DOSE:** Initial number of parasites fish was infected with
**DOB:** The date in YYYYMMDD the fish was born
**INFBIRTHDAYS:** Time in days between the first day of parental infection and fish birth
**PAIRBIRTHDAYS:** Time in days between the day the parental pair was set up to breed and the offspring's birth
**MAXWORMS:** The maximum number of worms recorded on the fish over its infection
**FULLAUC:** The area under the curve of infection load (number of worms) over time - 'infection integral'
**AGEATINF:** Age in days fish was when it was infected

The datasheet also contains the following variables about the infected parent. Where two variables are listed, those with an 'M' prefix refer to the mother, and those with an 'F' refer to the father:

**PARINFTREAT:** Which parent was infected - mother, father or neither.
**PINF:** Binary variable denoting that one parent was (1) or was not (0) infected.
**PAUC:** Parental (either the mother's or father's, depending on which was infected) area under the curve (AUC) of infection; NA if neither parent infected or parental AUC unknown
**MOTHERID/FATHERID:** ID of the parent
**MOTHERINF/FATHERINF:** Whether (1) or not (0) the parent was infected
**MMAXWORM/FMAXWORM:** The maximum number of worms recorded on the fish over its infection
**MAUC/FAUC:** The area under the curve of infection load (number of worms) over time - 'infection integral'
**MDATEINF/FDATEINF:** The date in YYYYMMDD the parent was infected.
**MPREW/FPREW:** Fish weight in mg before infection
**MPOSTW/FPOSTW:** Fish weight in mg after infection
**MPREL/FPREL:** Fish length in mm before infection
**MPOSTL/FPOSTL:** Fish length in mm after infection
**MpreSMI/FpreSMI:** Body condition - scaled mass index - before infection.
**MpostSMI/FpostSMI:** Body condition - scaled mass index - after infection.
**MdeltaSMI/FdeltaSMI:** Body condition - scaled mass index - change over the course of infection.
**MDIED/FDIED:** Whether or not the mother or father died during the observation period

### Data Visualization

We used pairwise plots to check for correlations between our variables of interest.

```r
## running pairs plot
panel.cor <- function(x, y, digits = 2, prefix = "", cex.cor,
    ...) {
    usr <- par("usr")
    on.exit(par(usr))
    par(usr = c(0, 1, 0, 1))
    r <- abs(cor(x, y))
    txt <- format(c(r, 0.123456789), digits = digits)[1]
    txt <- paste(prefix, txt, sep = "")
    if (missing(cex.cor))
        cex.cor <- 0.8/strwidth(txt)
    text(0.5, 0.5, txt, cex = cex.cor * r)
}

pairs(~FULLAUC + SEX + preSMI + LENGTHRESID + AGEATINF + INFBIRTHDAYS +
    PARINFTREAT + INFTYPE + PAIRBIRTHDAYS, data = IND, lower.panel = panel.smooth,
    upper.panel = panel.cor, na.action = na.omit)
```

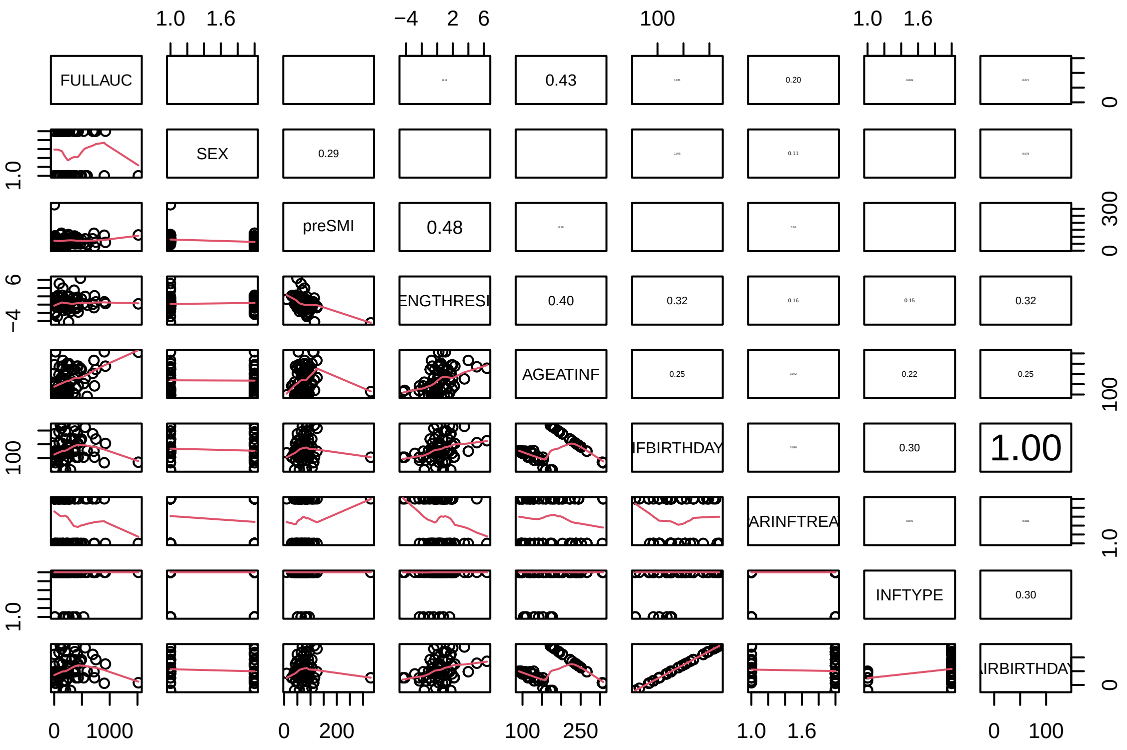

### Does infection affect female or male fitness?

#### Do infected and uninfected fish differ in fitness?

Parental infection increases latency to birth offspring, but infected and uninfected parents don't differ in the number, size, or body condition of offspring they produce. The result of m4 is presented in Figure 1 in the main text - infected mothers took longer to birth offspring than either pairs in which fathers had been infected, or both parents were naive to infection.

```r
# This question we're asking at the level of the individual
# mother

# Does infection affect number of babies, controlling for
# parental body condition and mother body length?
m1 <- glmmTMB(TOTBABIES ~ PARINFTREAT + MpostSMI + FpostSMI +
    MPREL, data = mot)
Anova(m1)
```

```
## Analysis of Deviance Table (Type II Wald chisquare tests)
##
## Response: TOTBABIES
##              Chisq Df Pr(>Chisq)
## PARINFTREAT 2.7207  2     0.2566
## MpostSMI    0.9518  1     0.3293
## FpostSMI    0.5284  1     0.4673
## MPREL       0.7120  1     0.3988
```

```r
sim_residuals_glmmTMB <- simulateResiduals(m1, 1000)
plot(sim_residuals_glmmTMB)
```

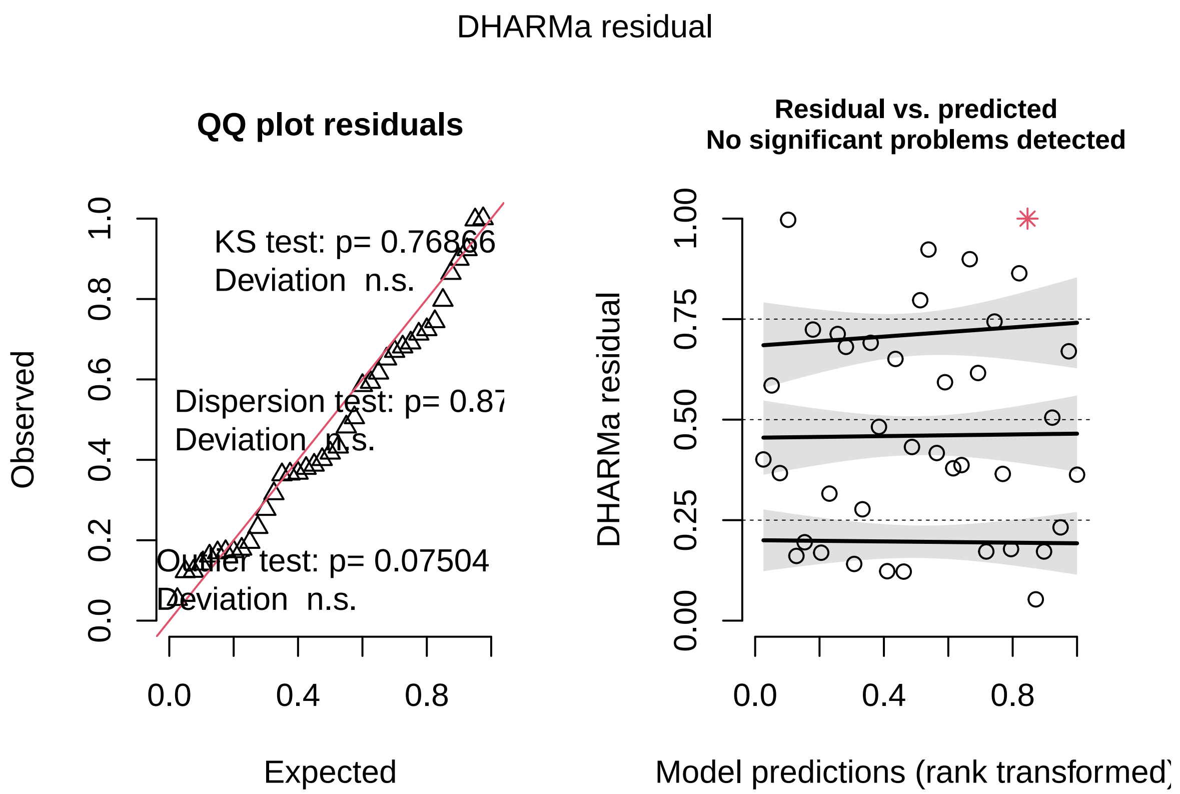

```
DHARMa::testDispersion(sim_residuals_glmmTMB)
```

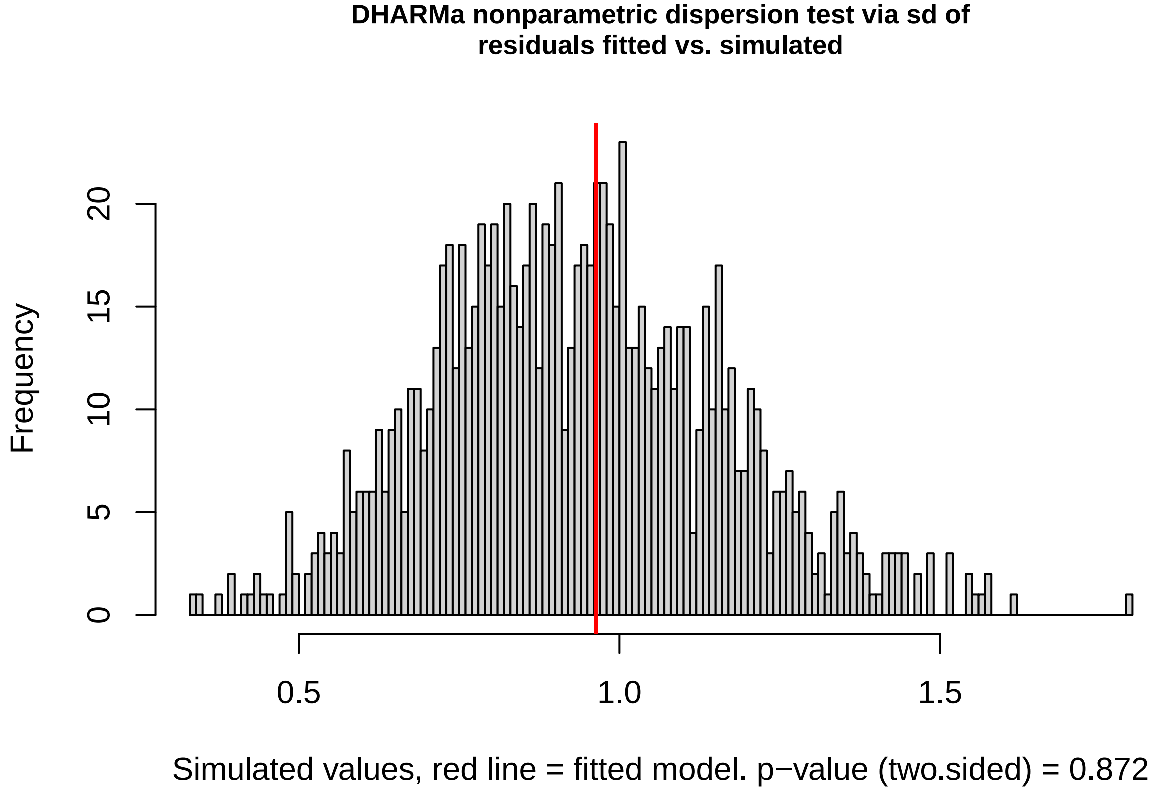

```
##
##  DHARMa nonparametric dispersion test via sd of residuals fitted vs.
```

```
##  simulated
##
## data:  simulationOutput
## dispersion = 1.0288, p-value = 0.872
## alternative hypothesis: two.sided
```

```r
# These questions we're asking at the level of the
# individual offspring

# Does infection affect the body condition at adulthood of
# offspring?
m2 <- glmmTMB(preSMI ~ PARINFTREAT + MdeltaSMI + FdeltaSMI +
    SEX + MPREL + (1 | MOTHERID), data = IND)
```

```
## Warning in finalizeTMB(TMBStruc, obj, fit, h, data.tmb.old): Model convergence
## problem; non-positive-definite Hessian matrix. See vignette('troubleshooting')
```

```r
Anova(m2)
```

```
## Analysis of Deviance Table (Type II Wald chisquare tests)
##
## Response: preSMI
##               Chisq Df Pr(>Chisq)
## PARINFTREAT  1.5191  2  0.4678873
## MdeltaSMI    1.5078  1  0.2194695
## FdeltaSMI    1.4294  1  0.2318571
## SEX         10.9386  1  0.0009418 ***
## MPREL        0.3980  1  0.5281432
## ---
## Signif. codes:  0 '***' 0.001 '**' 0.01 '*' 0.05 '.' 0.1 ' ' 1
```

```r
sim_residuals_glmmTMB <- simulateResiduals(m2, 1000)
plot(sim_residuals_glmmTMB)
```

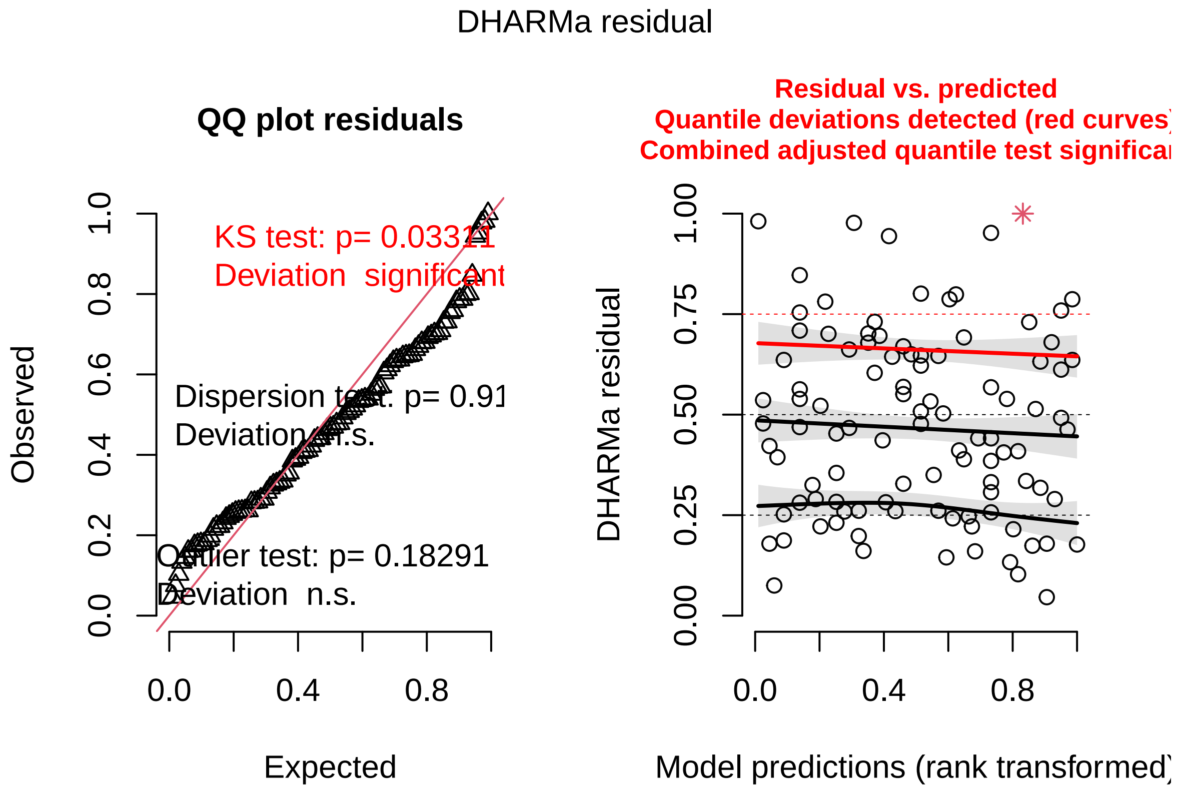

```
DHARMa::testDispersion(sim_residuals_glmmTMB)
```

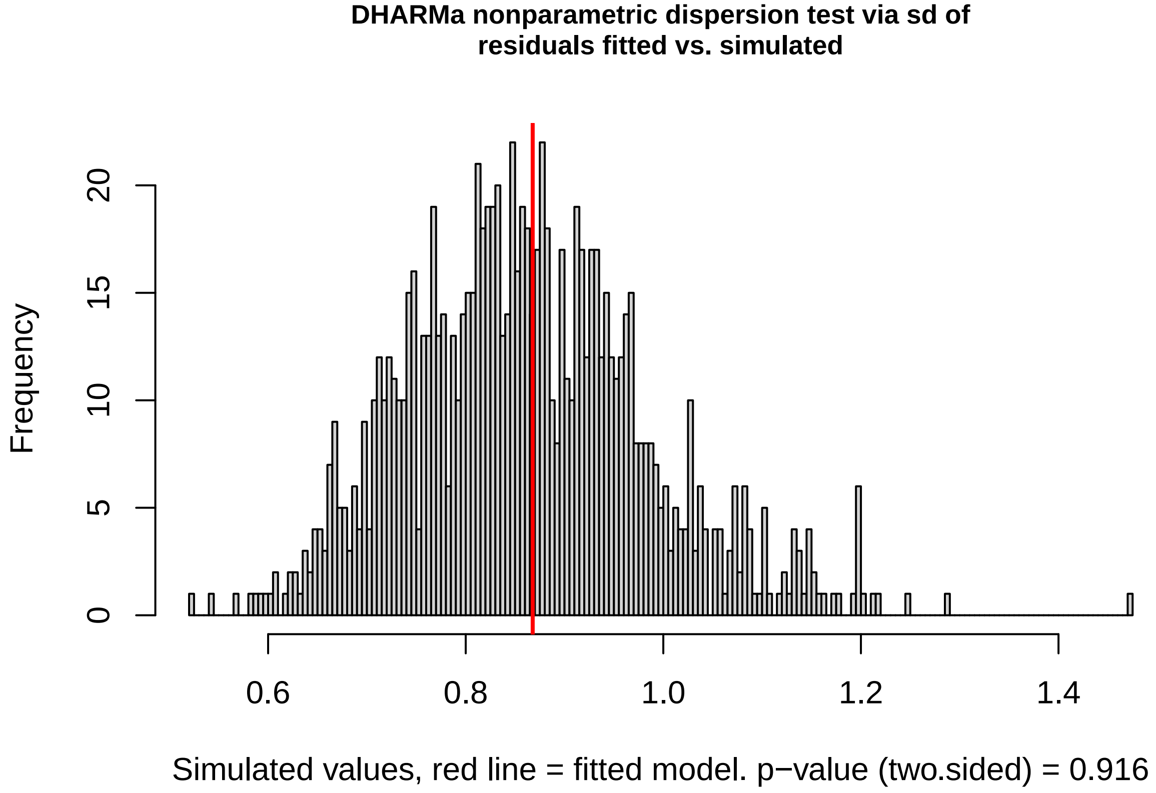

```
##
##  DHARMa nonparametric dispersion test via sd of residuals fitted vs.
```

```
##  simulated
##
## data:  simulationOutput
## dispersion = 1.0069, p-value = 0.916
## alternative hypothesis: two.sided
```

```
# Does infection affect the body length at adulthood of
# offspring?
m3 <- glmmTMB(LENGTHRESID ~ PARINFTREAT + MdeltaSMI + FdeltaSMI +
    SEX + MPREL + (1 | MOTHERID), data = IND)
Anova(m3)
```

```
## Analysis of Deviance Table (Type II Wald chisquare tests)
##
## Response: LENGTHRESID
##              Chisq Df Pr(>Chisq)
## PARINFTREAT 0.3323  2     0.8469
## MdeltaSMI   0.0878  1     0.7670
## FdeltaSMI   0.1532  1     0.6955
## SEX         0.1265  1     0.7221
## MPREL       0.7331  1     0.3919
```

```
sim_residuals_glmmTMB <- simulateResiduals(m3, 1000)
plot(sim_residuals_glmmTMB)
```

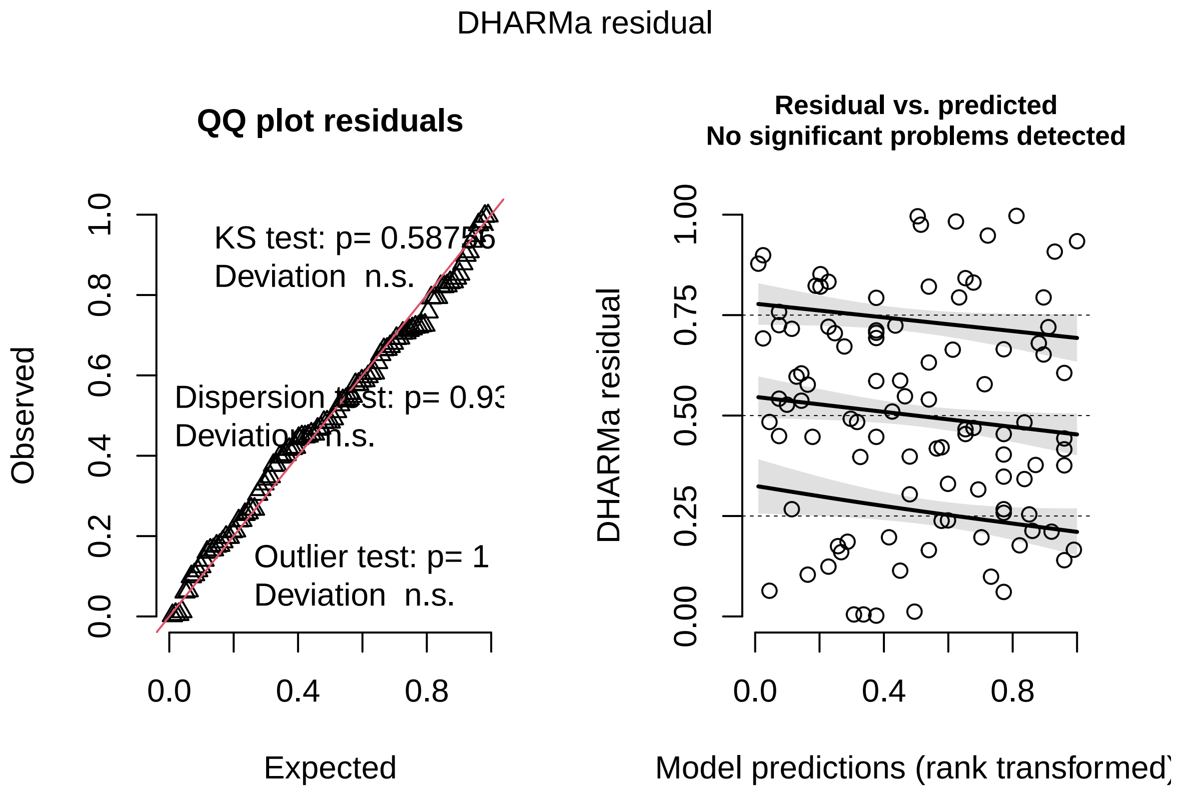

```
DHARMa::testDispersion(sim_residuals_glmmTMB)
```

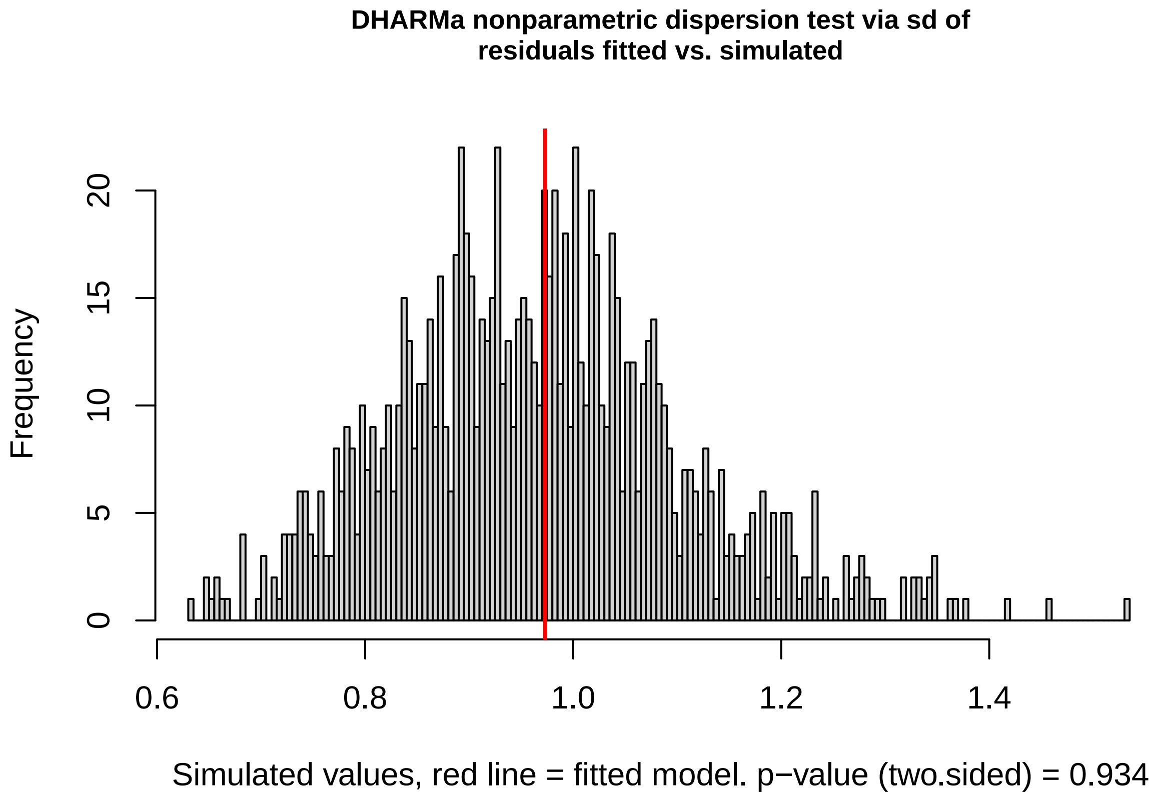

```
##
##  DHARMa nonparametric dispersion test via sd of residuals fitted vs.
##  simulated
##
## data:  simulationOutput
## dispersion = 1.0066, p-value = 0.934
## alternative hypothesis: two.sided
```

```r
# Does infection affect the latency to give birth?
m4 <- glmmTMB(PAIRBIRTHDAYS ~ PARINFTREAT + MpostSMI + FpostSMI +
    MPREL + (1 | MOTHERID), data = IND)
Anova(m4)
```

```
## Analysis of Deviance Table (Type II Wald chisquare tests)
##
## Response: PAIRBIRTHDAYS
##              Chisq Df Pr(>Chisq)
## PARINFTREAT 0.5969  2     0.7420
## MpostSMI    0.0649  1     0.7988
## FpostSMI    1.9265  1     0.1651
## MPREL       0.0000  1     0.9947
```

```r
sim_residuals_glmmTMB <- simulateResiduals(m4, 1000)
plot(sim_residuals_glmmTMB)
```

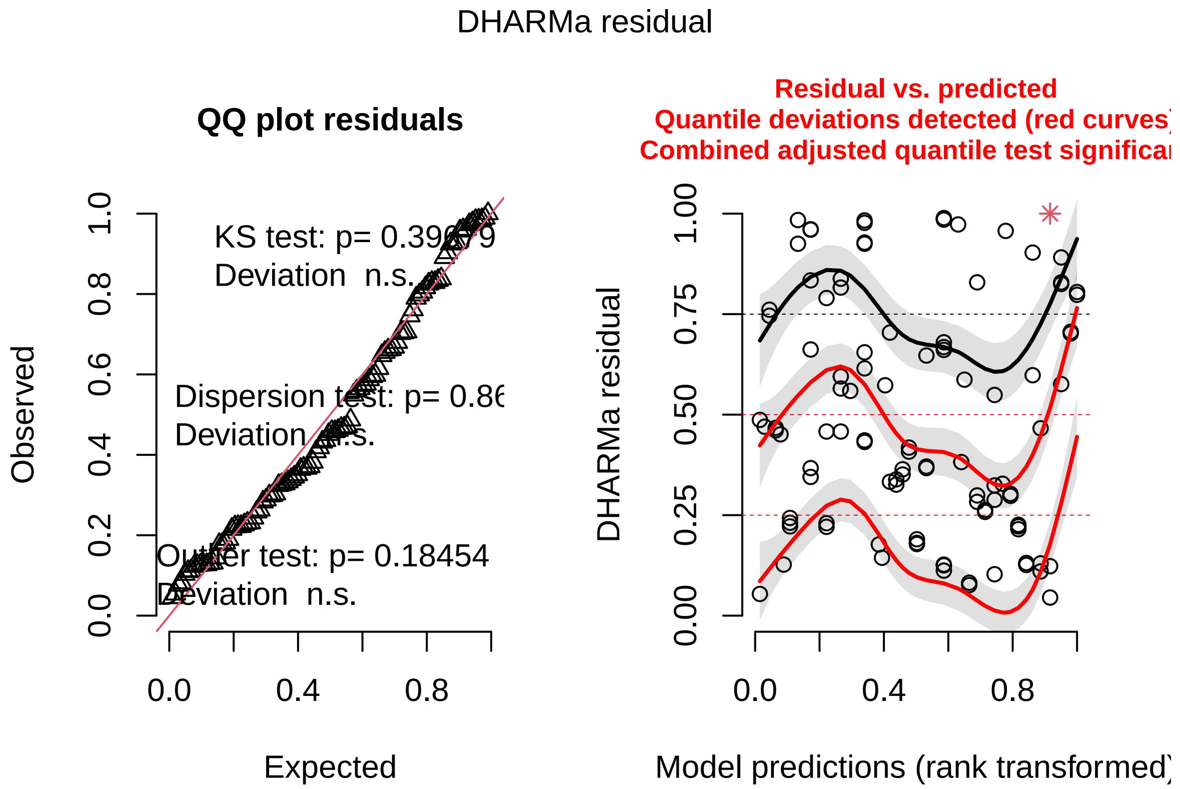

```r
DHARMa::testDispersion(sim_residuals_glmmTMB)
```

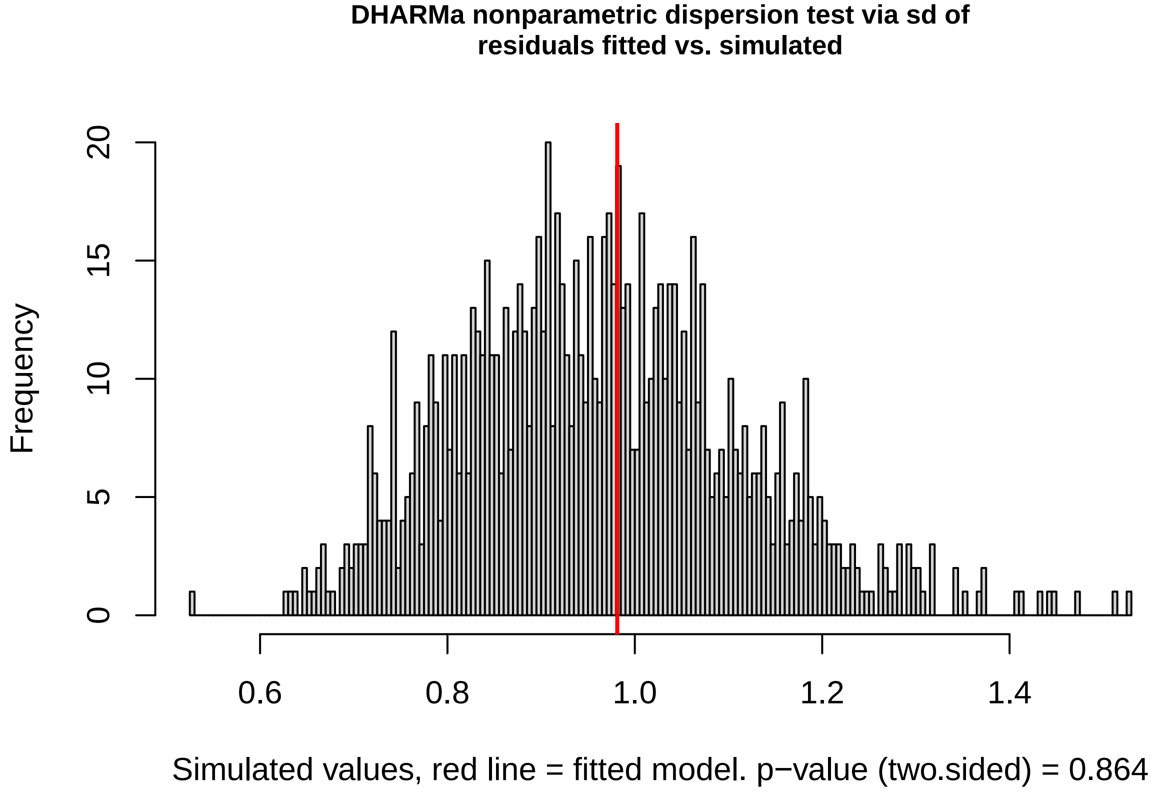

```
##
##  DHARMa nonparametric dispersion test via sd of residuals fitted vs.
```

```
##  simulated
##
## data:  simulationOutput
## dispersion = 1.0188, p-value = 0.864
## alternative hypothesis: two.sided
```

##### Among infected parents, does infection severity correlate with fitness?

Here we subset the analysis to look at just pairs where one parent was infected to ask whether the severity of that infection predicted the number, size, or body condition of offspring, and the latency to birth. The result of this model is presented in Figure 1 in the main text.

```
# This question we're asking at the level of the individual
# mother

# Does infection severity among infected parents affect
# number of babies, controlling for parental body condition
# and mother body length?
m5 <- glmmTMB(TOTBABIES ~ log(PAUC + 1) * PARINFTREAT + MpostSMI +
    FpostSMI + MPREL, data = motinf)
Anova(m5)
```

```
## Analysis of Deviance Table (Type II Wald chisquare tests)
##
## Response: TOTBABIES
##                            Chisq Df Pr(>Chisq)
## log(PAUC + 1)             0.0004  1     0.9831
## PARINFTREAT               2.3809  1     0.1228
## MpostSMI                  1.4531  1     0.2280
## FpostSMI                  0.2347  1     0.6280
## MPREL                     0.9940  1     0.3188
## log(PAUC + 1):PARINFTREAT 0.0684  1     0.7937
```

```
sim_residuals_glmmTMB <- simulateResiduals(m5, 1000)
plot(sim_residuals_glmmTMB)
```

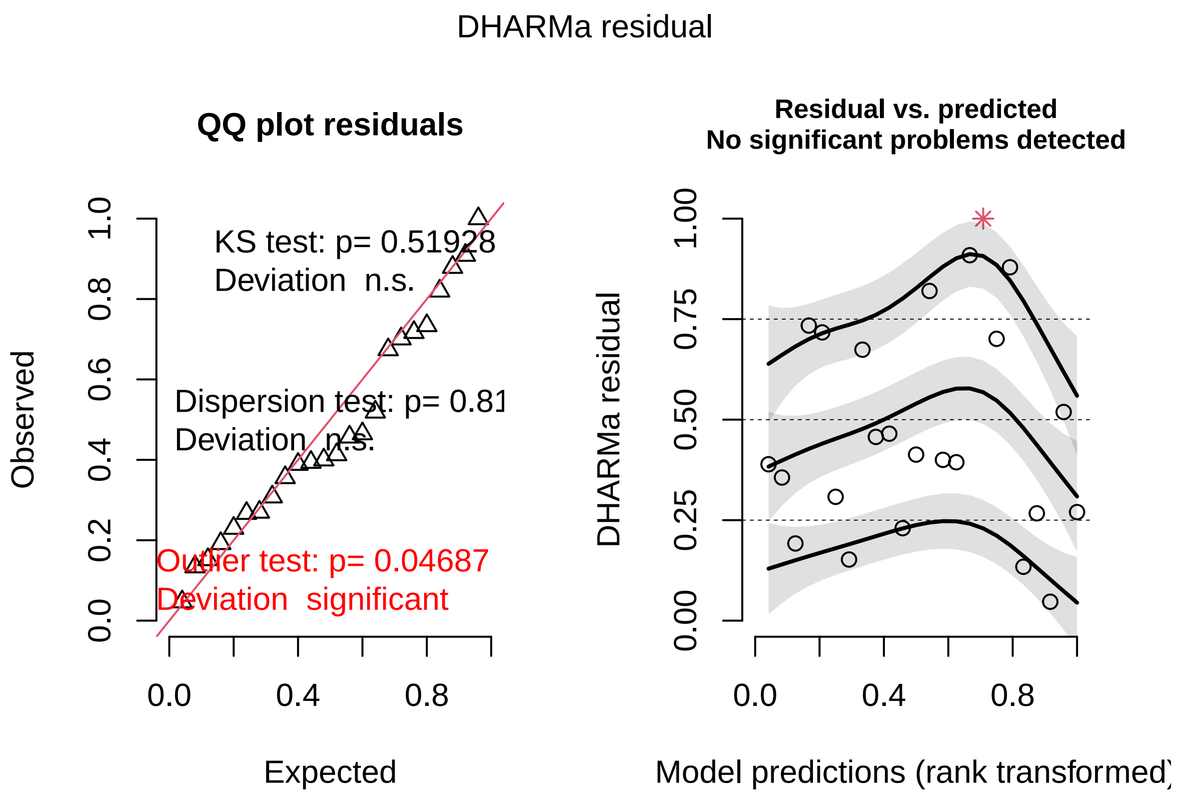

```
DHARMa::testDispersion(sim_residuals_glmmTMB)
```

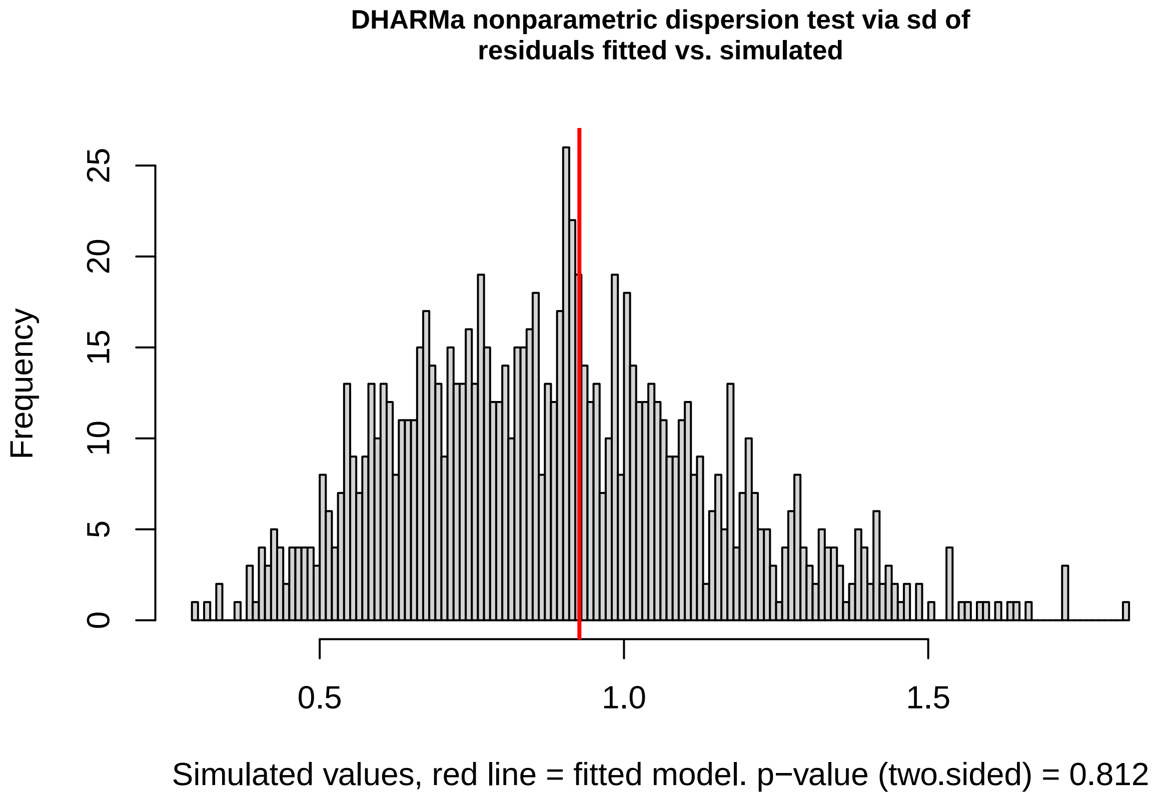

```
##
##  DHARMa nonparametric dispersion test via sd of residuals fitted vs.
```

```
##  simulated
##
## data:  simulationOutput
## dispersion = 1.0424, p-value = 0.812
## alternative hypothesis: two.sided
```

```
# These questions we're asking at the level of the
# individual offspring

m6 <- glmmTMB(preSMI ~ log(PAUC + 1) * PARINFTREAT + MpostSMI +
    FpostSMI + SEX + (1 | MOTHERID), data = INDPARINFECTED)
Anova(m6)
```

```
## Analysis of Deviance Table (Type II Wald chisquare tests)
##
## Response: preSMI
##                            Chisq Df Pr(>Chisq)
## log(PAUC + 1)             3.5134  1   0.060875 .
## PARINFTREAT               0.0020  1   0.964683
## MpostSMI                  0.2917  1   0.589153
## FpostSMI                  0.0897  1   0.764506
## SEX                       6.6514  1   0.009908 **
## log(PAUC + 1):PARINFTREAT 1.1180  1   0.290341
## ---
## Signif. codes:  0 '***' 0.001 '**' 0.01 '*' 0.05 '.' 0.1 ' ' 1
```

```
sim_residuals_glmmTMB <- simulateResiduals(m6, 1000)
plot(sim_residuals_glmmTMB)
```

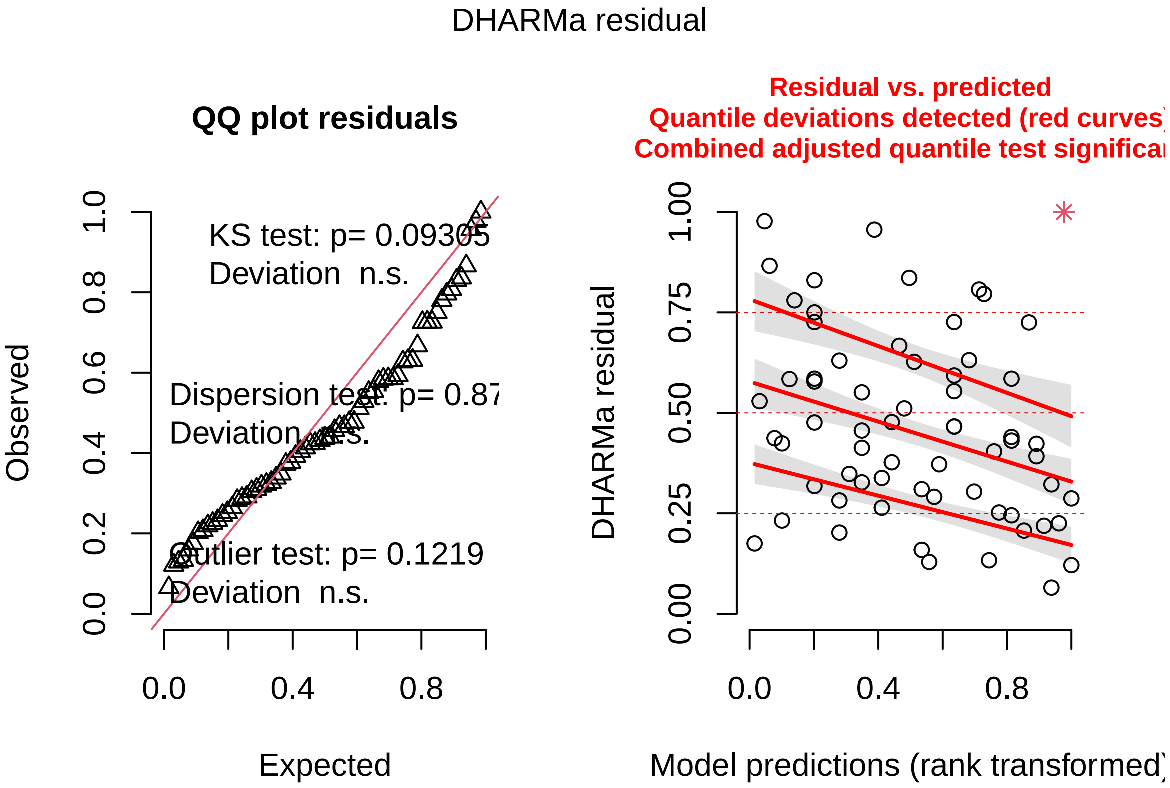

```
DHARMa::testDispersion(sim_residuals_glmmTMB)
```

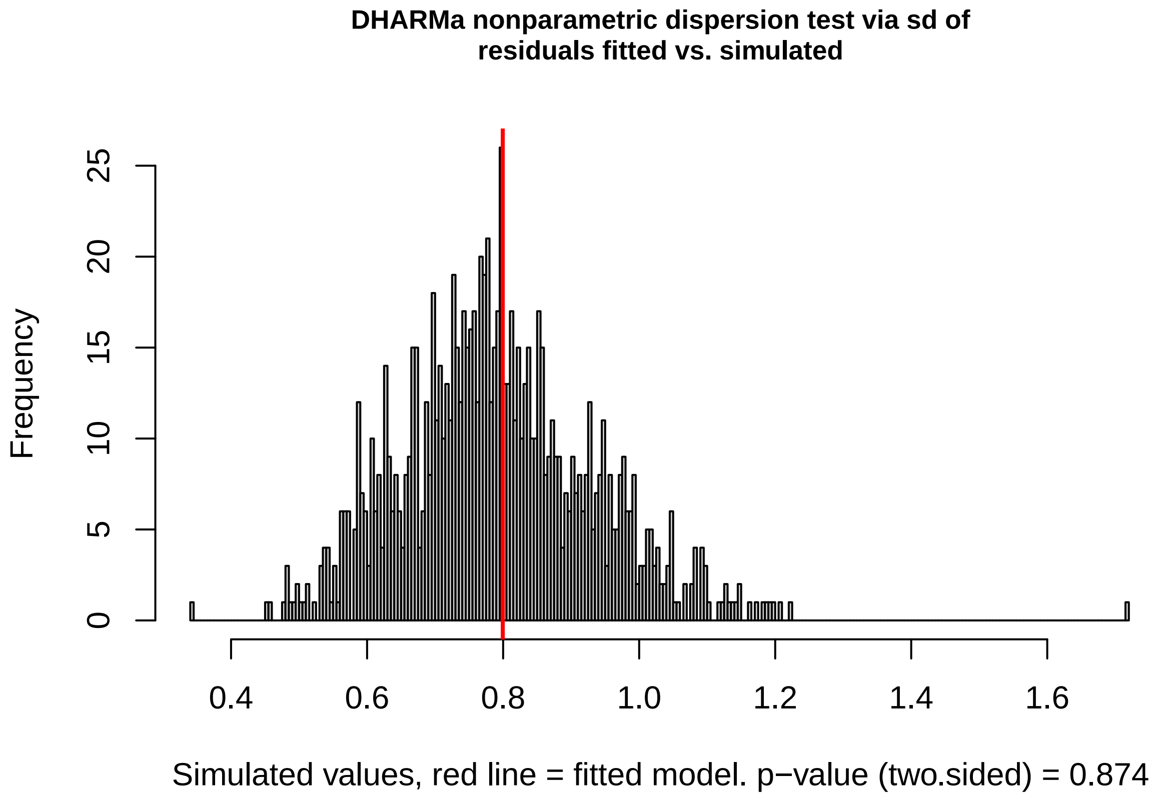

```
##
##  DHARMa nonparametric dispersion test via sd of residuals fitted vs.
##  simulated
##
## data:  simulationOutput
## dispersion = 1.0099, p-value = 0.874
## alternative hypothesis: two.sided
```

```
m7 <- glmmTMB(LENGTHRESID ~ log(PAUC + 1) + PARINFTREAT + MpostSMI +
    FpostSMI + SEX + MPREL + (1 | MOTHERID), data = INDPARINFECTED)
Anova(m7)
```

```
## Analysis of Deviance Table (Type II Wald chisquare tests)
##
## Response: LENGTHRESID
##                Chisq Df Pr(>Chisq)
## log(PAUC + 1) 0.0010  1     0.9745
## PARINFTREAT   0.6579  1     0.4173
## MpostSMI      0.3759  1     0.5398
## FpostSMI      1.1640  1     0.2806
## SEX           0.0247  1     0.8751
## MPREL         0.3276  1     0.5671
```

```
sim_residuals_glmmTMB <- simulateResiduals(m7, 1000)
plot(sim_residuals_glmmTMB)
```

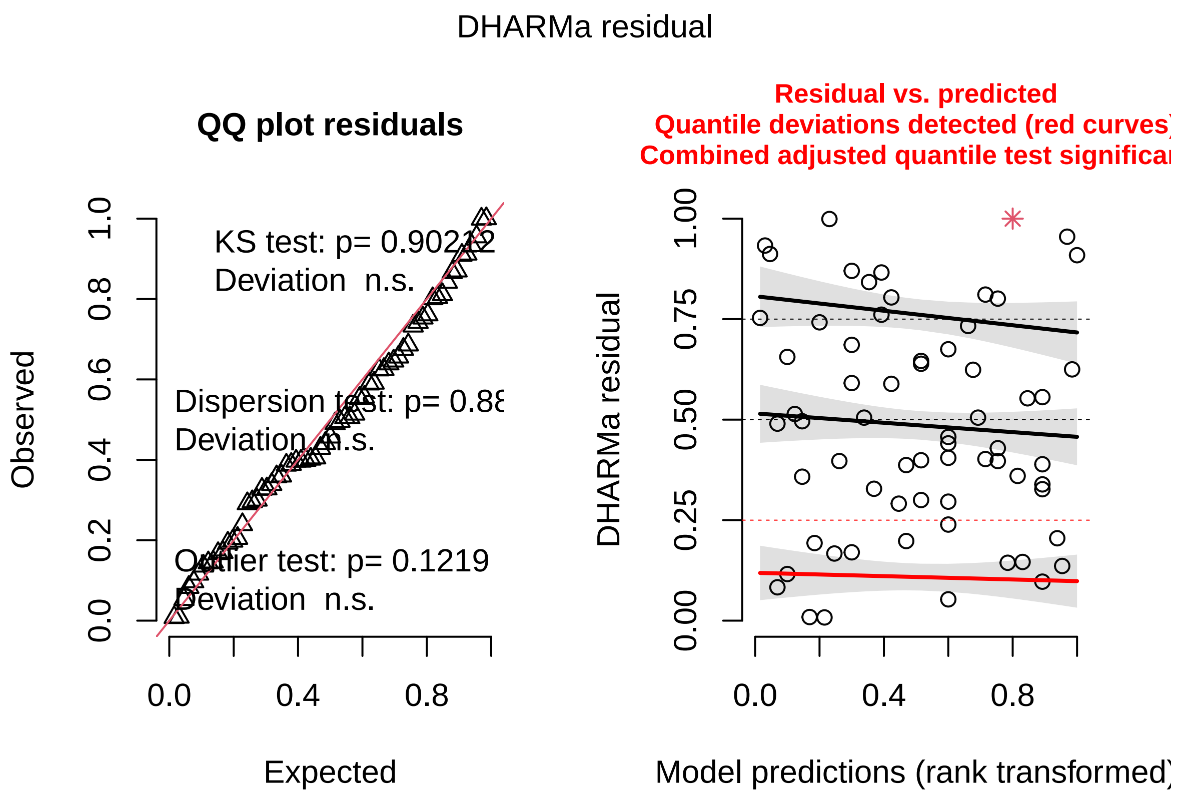

```
DHARMa::testDispersion(sim_residuals_glmmTMB)
```

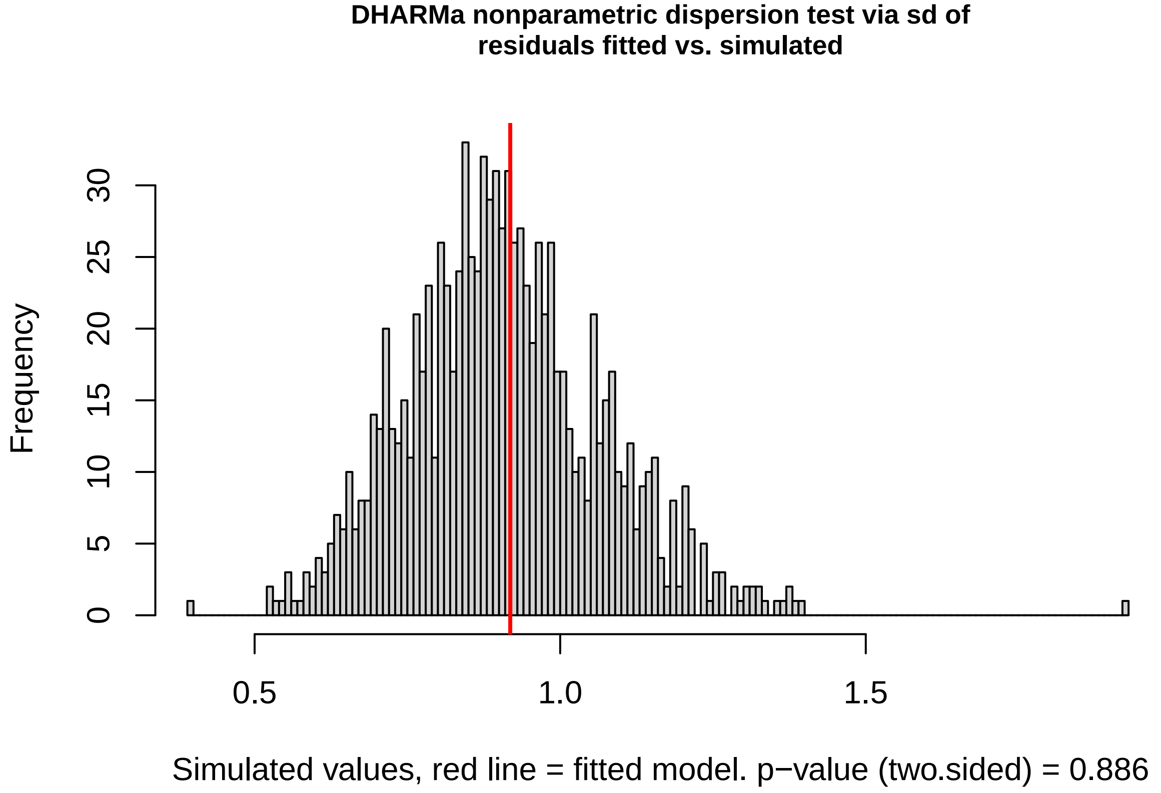

```
##
##  DHARMa nonparametric dispersion test via sd of residuals fitted vs.
```

```
##  simulated
##
## data:  simulationOutput
## dispersion = 1.0113, p-value = 0.886
## alternative hypothesis: two.sided
```

```
m8 <- glmmTMB(INFBIRTHDAYS ~ log(PAUC + 1) * PARINFTREAT + MpostSMI +
    FpostSMI + MPREL + (1 | MOTHERID), data = INDPARINFECTED)
Anova(m8)
```

```
## Analysis of Deviance Table (Type II Wald chisquare tests)
##
## Response: INFBIRTHDAYS
##                              Chisq Df Pr(>Chisq)
## log(PAUC + 1)               0.2192  1    0.63963
## PARINFTREAT                 0.0097  1    0.92156
## MpostSMI                    0.4500  1    0.50236
## FpostSMI                    0.0331  1    0.85567
## MPREL                       0.2116  1    0.64549
## log(PAUC + 1):PARINFTREAT 5.1294  1    0.02352 *
## ---
## Signif. codes:  0 '***' 0.001 '**' 0.01 '*' 0.05 '.' 0.1 ' ' 1
```

```
sim_residuals_glmmTMB <- simulateResiduals(m8, 1000)
plot(sim_residuals_glmmTMB)
```

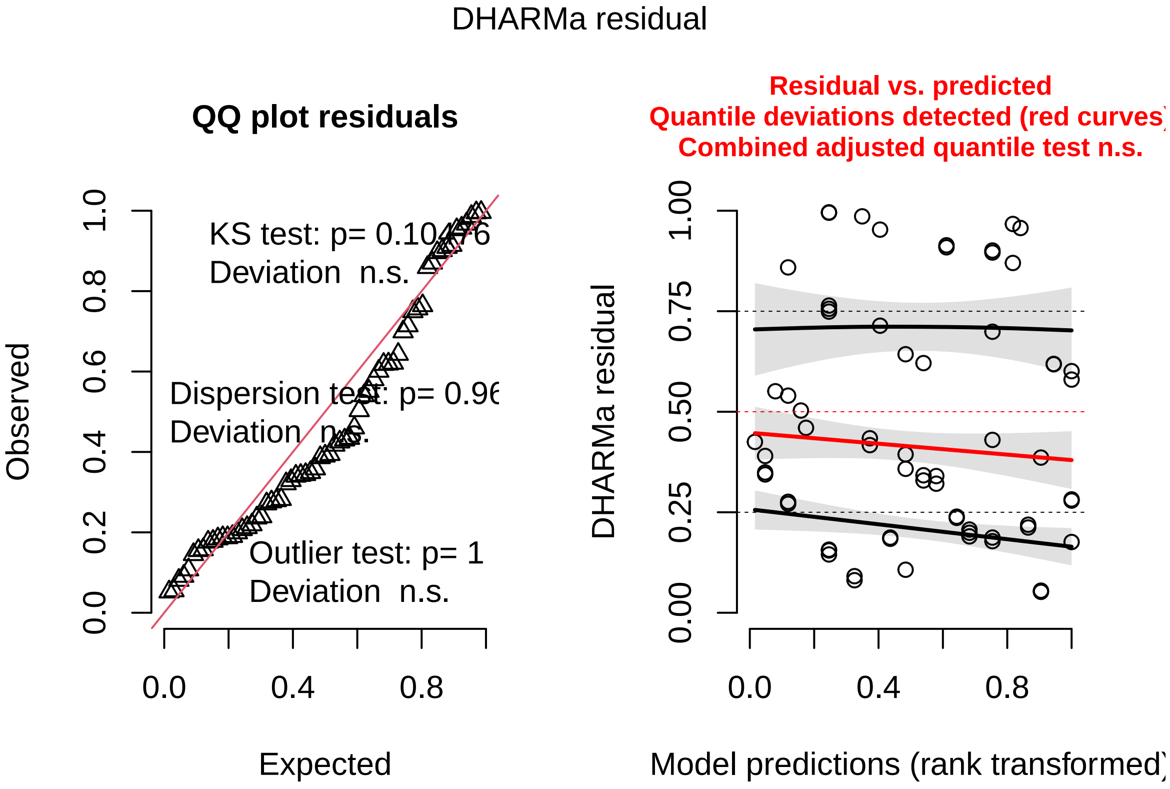

```
DHARMa::testDispersion(sim_residuals_glmmTMB)
```

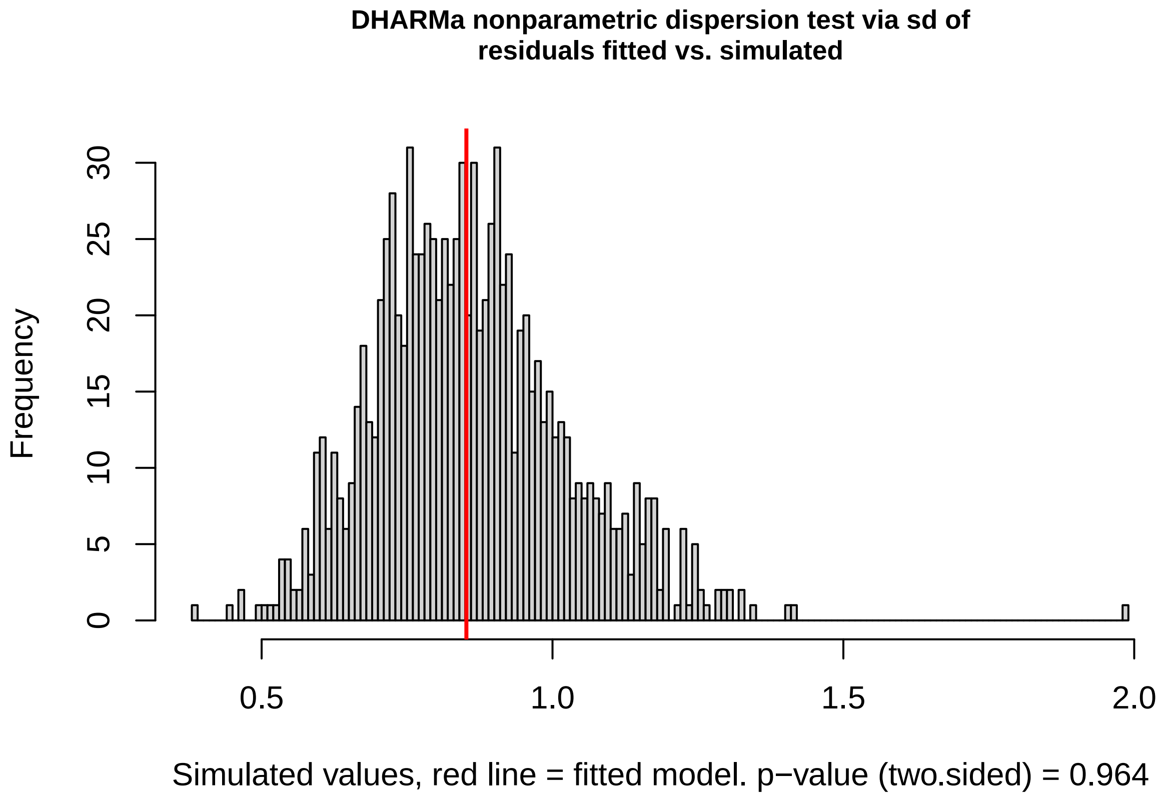

```
##
##  DHARMa nonparametric dispersion test via sd of residuals fitted vs.
##  simulated
##
## data:  simulationOutput
## dispersion = 0.99113, p-value = 0.964
## alternative hypothesis: two.sided
```

##### Is the effect of parental infection severity on birth latency significantly different from 0, or do the sexes just differ?

The correlation in Figure 1 among infected mothers is not significantly different from 0, where as that among infected fathers is significantly different from 0.

```
# Subset INDPARINFECTED for infected fathers
INDPARINFECTEDFATHER <- subset(INDPARINFECTED, PARINFTREAT ==
    "FATHER")
# Subset INDPARINFECTED for infected mothers
INDPARINFECTEDMOTHER <- subset(INDPARINFECTED, PARINFTREAT ==
    "MOTHER")
```

```
m8m <- glmmTMB(INFBIRTHDAYS ~ log(PAUC + 1) + (1 | MOTHERID),
    data = INDPARINFECTEDMOTHER)
Anova(m8m)
```

```
## Analysis of Deviance Table (Type II Wald chisquare tests)
##
## Response: INFBIRTHDAYS
##                 Chisq Df Pr(>Chisq)
## log(PAUC + 1) 0.8721  1     0.3504
```

```
sim_residuals_glmmTMB <- simulateResiduals(m8m, 1000)
plot(sim_residuals_glmmTMB)
```

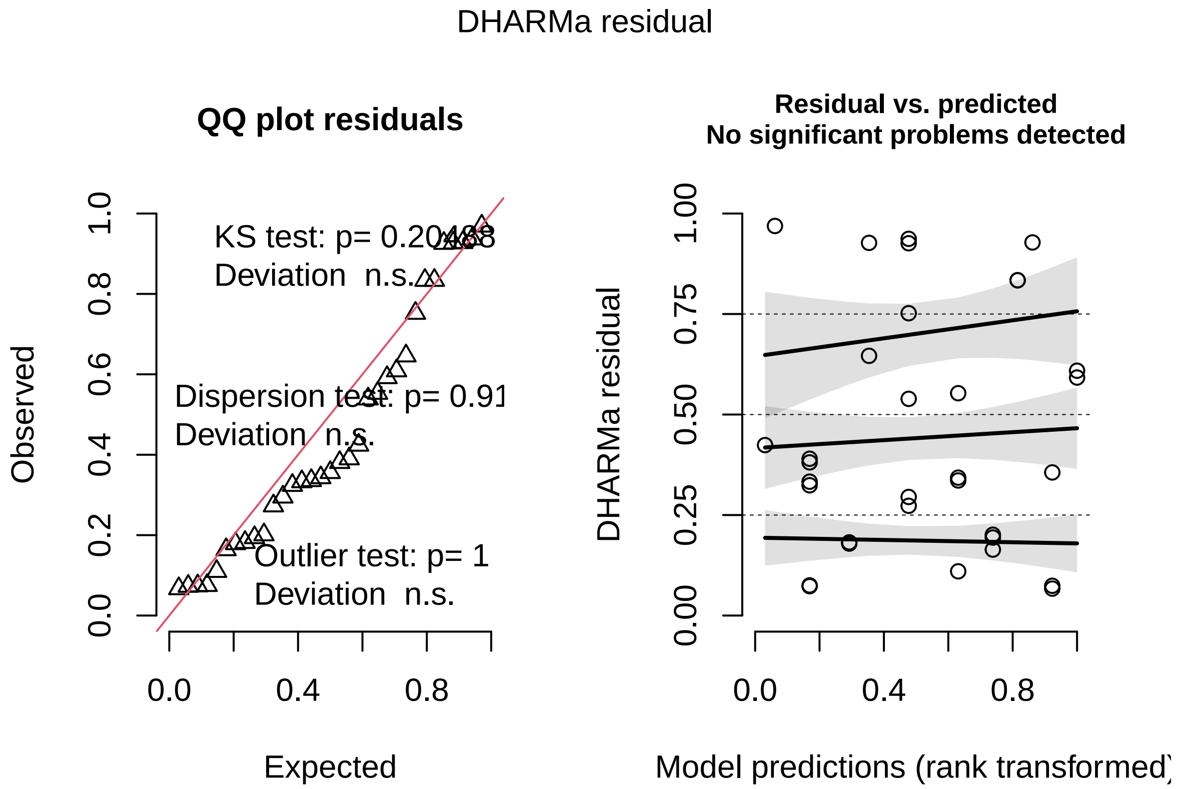

```
DHARMa::testDispersion(sim_residuals_glmmTMB)
```

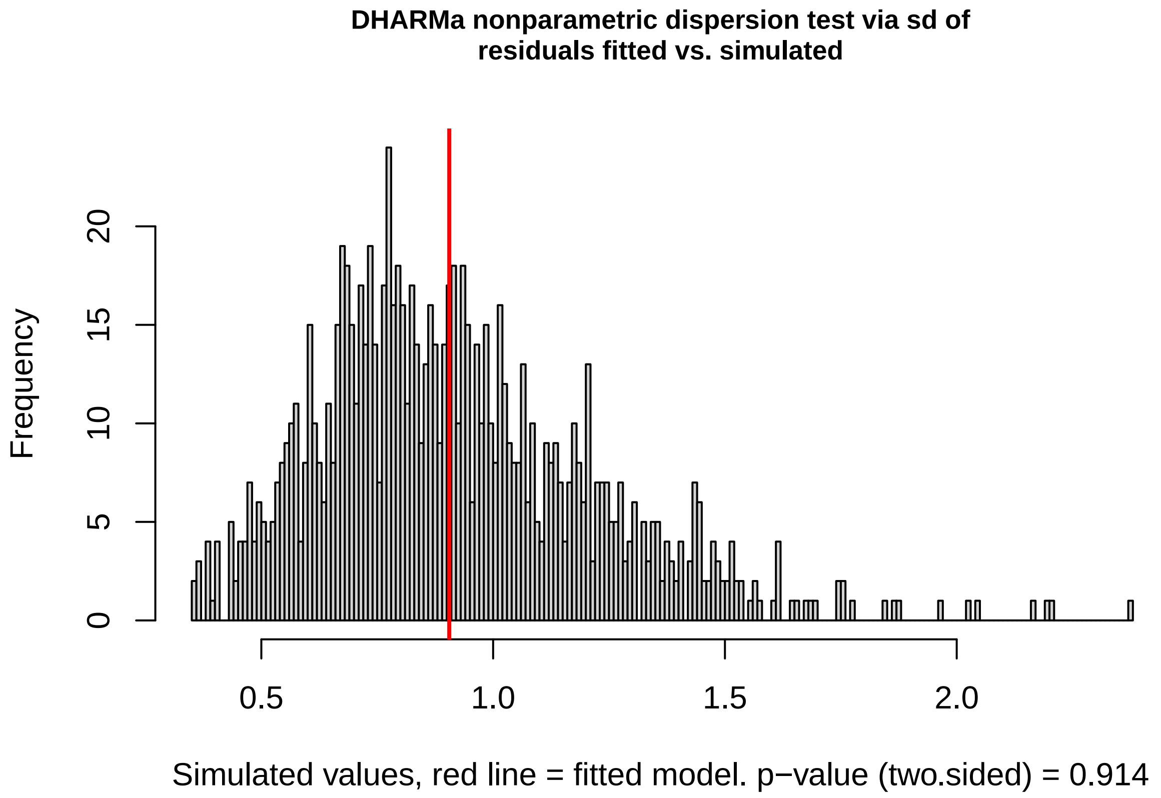

```
##
##  DHARMa nonparametric dispersion test via sd of residuals fitted vs.
##  simulated
##
## data:  simulationOutput
## dispersion = 0.98563, p-value = 0.914
## alternative hypothesis: two.sided
```

```
m8f <- glmmTMB(INFBIRTHDAYS ~ log(PAUC + 1) + (1 | MOTHERID),
    data = INDPARINFECTEDFATHER)
Anova(m8f)
```

```
## Analysis of Deviance Table (Type II Wald chisquare tests)
##
## Response: INFBIRTHDAYS
##                Chisq Df Pr(>Chisq)
## log(PAUC + 1) 5.4989  1    0.01903 *
## ---
## Signif. codes:  0 '***' 0.001 '**' 0.01 '*' 0.05 '.' 0.1 ' ' 1
```

```
sim_residuals_glmmTMB <- simulateResiduals(m8f, 1000)
plot(sim_residuals_glmmTMB)
```

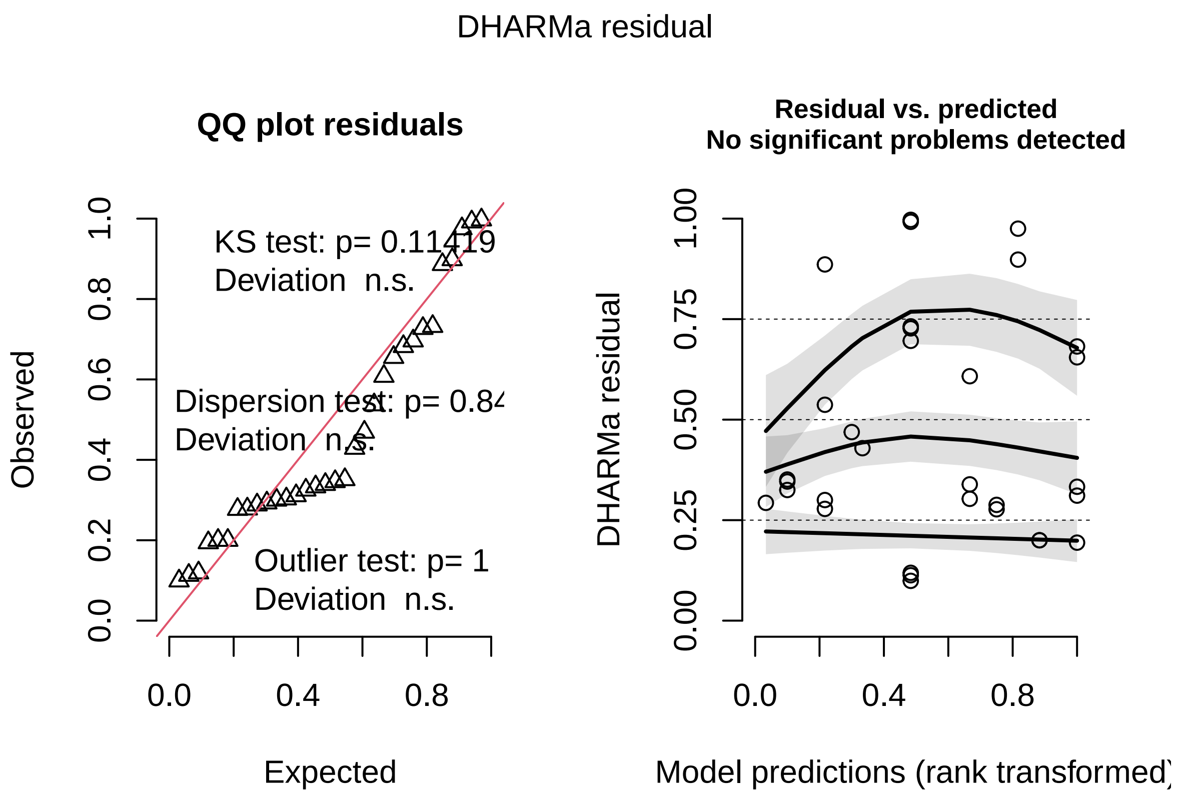

```
DHARMa::testDispersion(sim_residuals_glmmTMB)
```

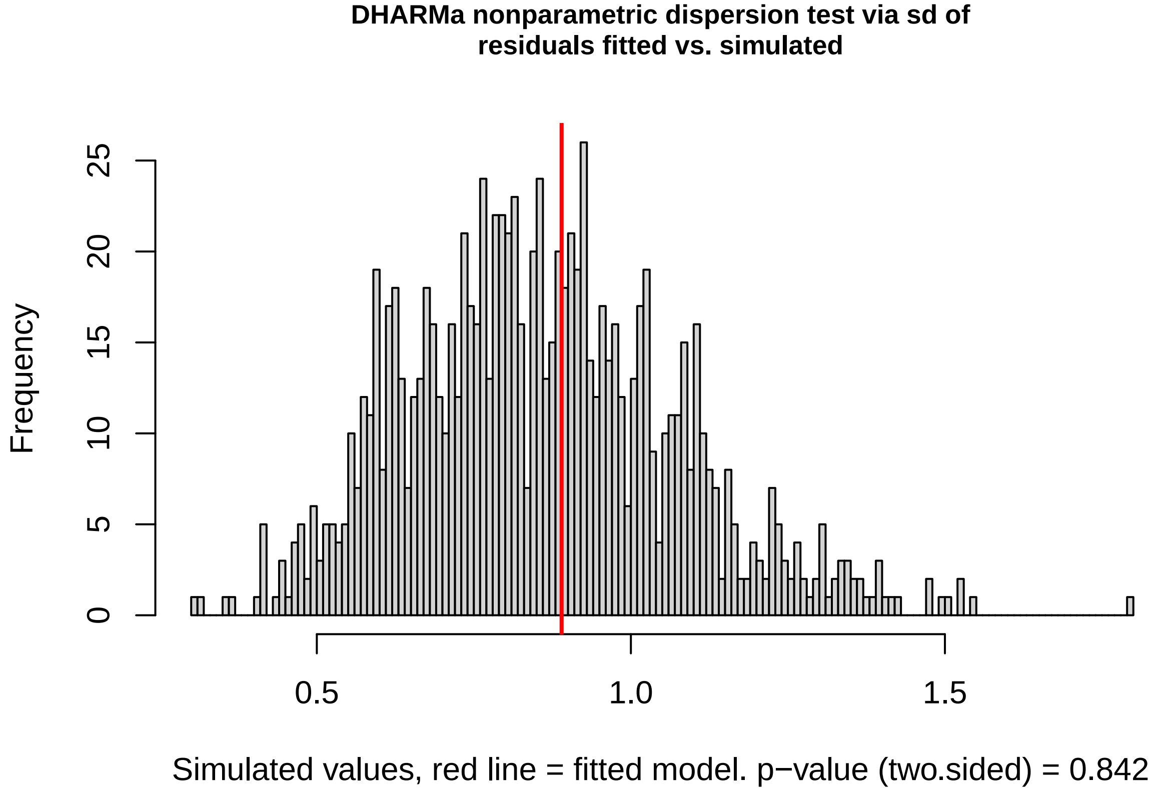

```
##
##  DHARMa nonparametric dispersion test via sd of residuals fitted vs.
```

```
##  simulated
##
## data:  simulationOutput
## dispersion = 1.038, p-value = 0.842
## alternative hypothesis: two.sided
```

### Does parental infection affect offspring fitness?

#### Do offspring of infected parents differ in their response to infection, compared to offspring of uninfected parents?

In this analysis, we tested whether the offspring of pairs in which one parent was infected differed in their response to infection from offspring of parasite-naive pairs.

```
m9 <- glmmTMB(FULLAUC ~ SEX + preSMI + PARINFTREAT + LENGTHRESID +
    DOSE + INFTYPE + AGEATINF + AGEATINF:PARINFTREAT + SEX:PARINFTREAT +
    (1 | MOTHERID), data = IND, na.action = na.omit)

Anova(m9)
```

```
## Analysis of Deviance Table (Type II Wald chisquare tests)
##
## Response: FULLAUC
##                       Chisq Df Pr(>Chisq)
## SEX                  0.0805  1   0.776590
## preSMI               0.0621  1   0.803257
## PARINFTREAT          2.2909  2   0.318074
## LENGTHRESID          0.0449  1   0.832187
## DOSE                 9.6880  1   0.001855 **
## INFTYPE              2.5991  1   0.106927
## AGEATINF             7.7161  1   0.005473 **
## PARINFTREAT:AGEATINF 7.3866  2   0.024890 *
## SEX:PARINFTREAT      0.4933  2   0.781407
## ---
## Signif. codes:  0 '***' 0.001 '**' 0.01 '*' 0.05 '.' 0.1 ' ' 1
```

```
sim_residuals_glmmTMB <- simulateResiduals(m9, 1000)
plot(sim_residuals_glmmTMB)
```

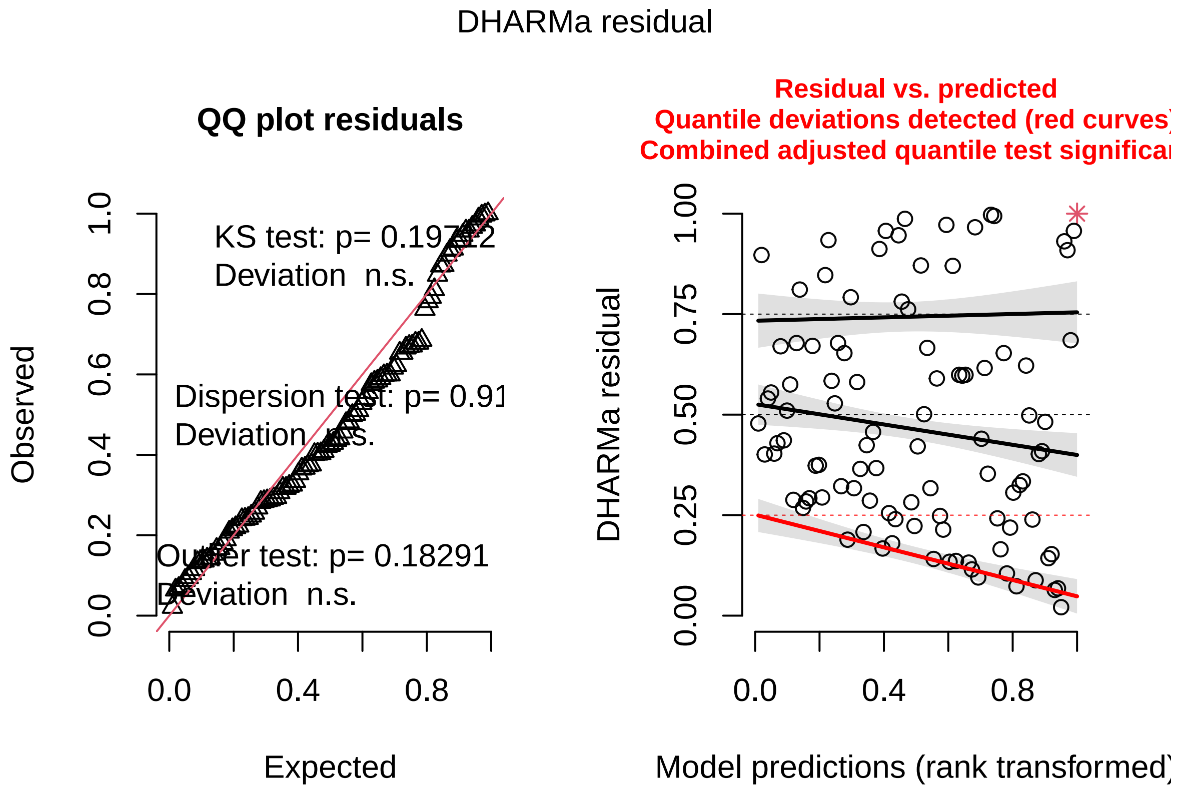

```
DHARMa::testDispersion(sim_residuals_glmmTMB)
```

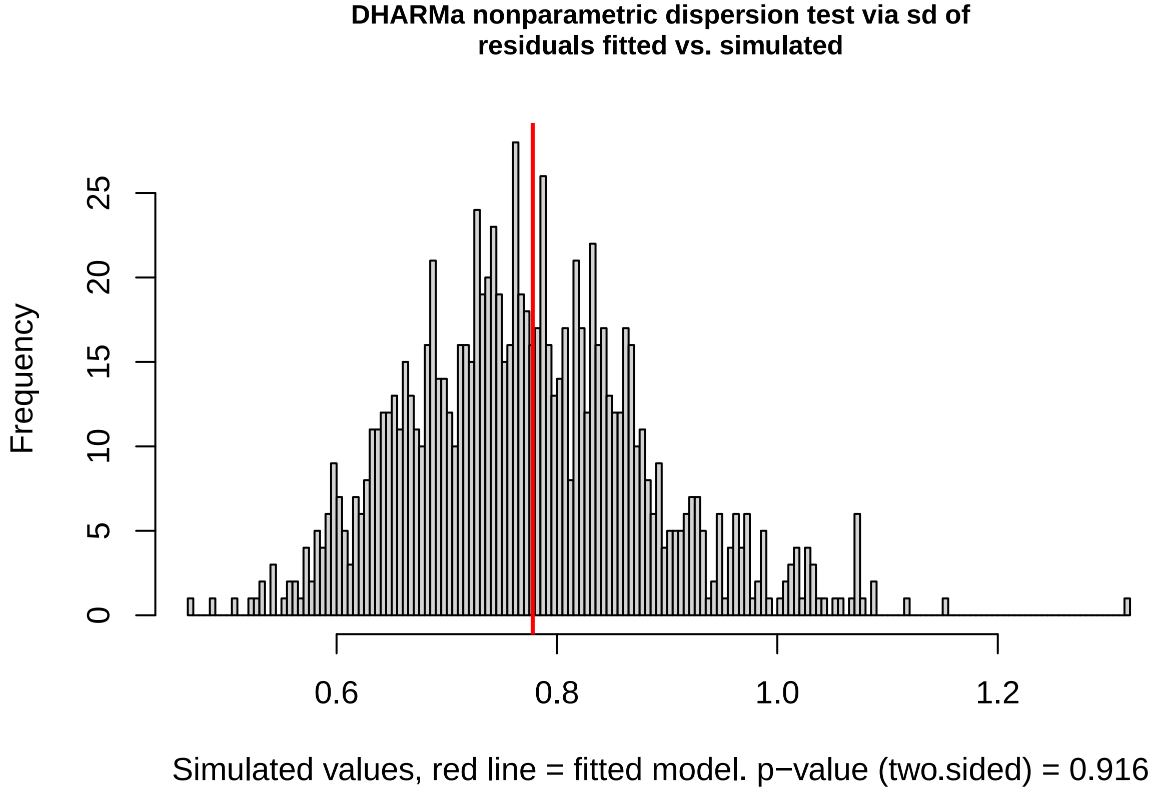

```
##
##  DHARMa nonparametric dispersion test via sd of residuals fitted vs.
```

```
##  simulated
##
## data:  simulationOutput
## dispersion = 1.0069, p-value = 0.916
## alternative hypothesis: two.sided
```

#### Is the correlation between age and offspring infection integral different from 0 among all parental treatment groups?

Below we run posthoc tests to evaluate whether the correlation in Fig. 2 differ from 0 for the "NEITHER" parental infection group. The other groups are evaluated in the posthoc tests below, for Model 10 (m10). The correlation here is not significantly different from 0.

```
INDn <- subset(IND, PARINFTREAT == "NEITHER")

m9n <- glmmTMB(FULLAUC ~ SEX + preSMI + LENGTHRESID + DOSE +
    INFTYPE + AGEATINF + (1 | MOTHERID), data = INDn, na.action = na.omit)
Anova(m9n)
```

```
## Analysis of Deviance Table (Type II Wald chisquare tests)
##
## Response: FULLAUC
##              Chisq Df Pr(>Chisq)
## SEX         1.0594  1    0.30336
## preSMI      2.8272  1    0.09268 .
## LENGTHRESID 2.7082  1    0.09983 .
## DOSE        6.1633  1    0.01304 *
## INFTYPE     1.5282  1    0.21638
## AGEATINF    2.3870  1    0.12235
## ---
## Signif. codes:  0 '***' 0.001 '**' 0.01 '*' 0.05 '.' 0.1 ' ' 1
```

```
sim_residuals_glmmTMB <- simulateResiduals(m9n, 1000)
plot(sim_residuals_glmmTMB)
```

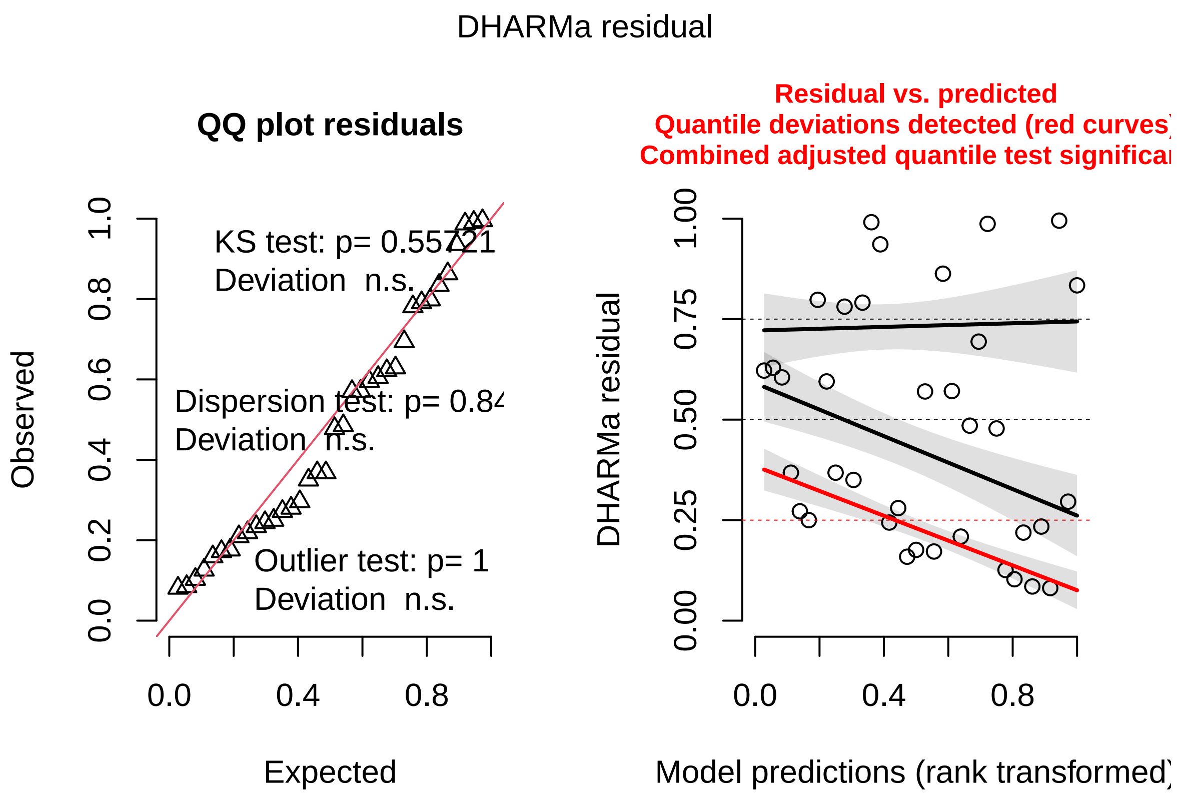

```
DHARMa::testDispersion(sim_residuals_glmmTMB)
```

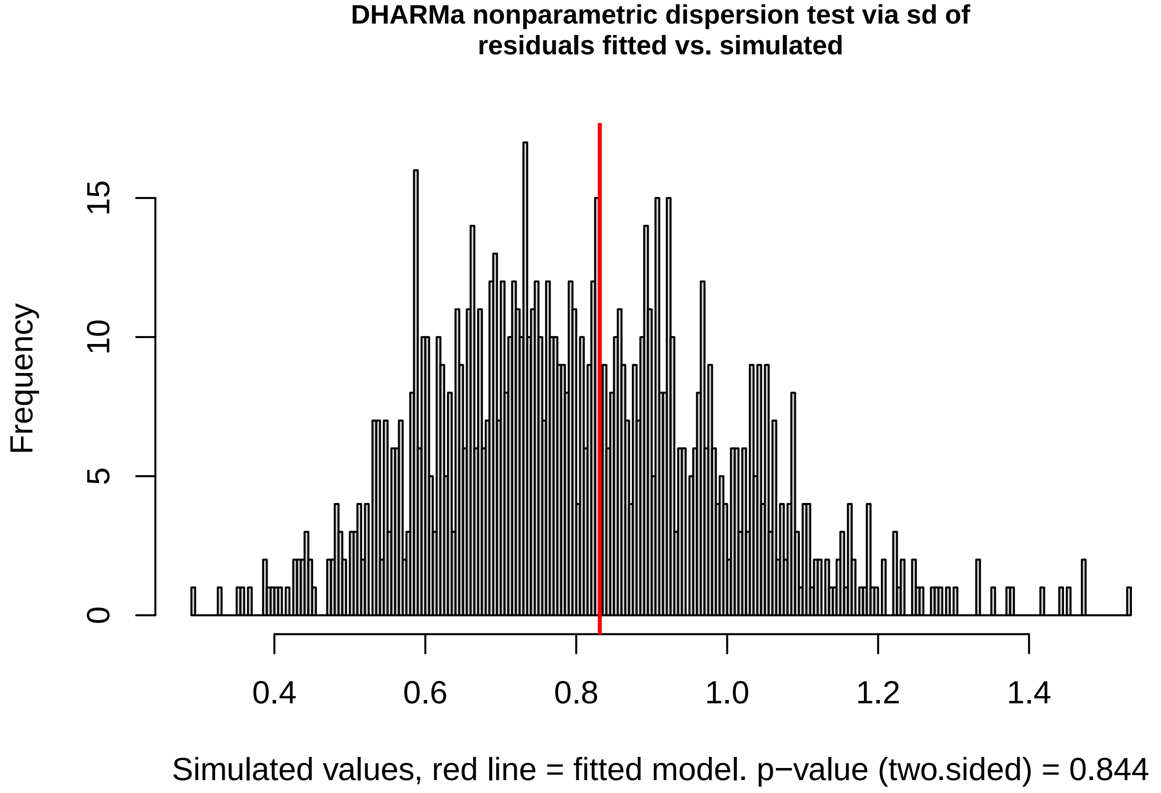

```
##
##  DHARMa nonparametric dispersion test via sd of residuals fitted vs.
```

```
##  simulated
##
## data:  simulationOutput
## dispersion = 1.0307, p-value = 0.844
## alternative hypothesis: two.sided
```

#### Among offspring with one infected parent, does parent infection predict offspring infection?

Here we used data only from offspring with one infected parent.

```
m10 <- glmmTMB(FULLAUC ~ log(PAUC + 1) * PARINFTREAT + log(PAUC +
    1):PARINFTREAT:SEX + SEX + preSMI + LENGTHRESID + DOSE +
    INFTYPE + AGEATINF + AGEATINF:PARINFTREAT + (1 | MOTHERID),
    data = INDPARINFECTED, na.action = na.omit, family = gaussian())

Anova(m10)
```

```
## Analysis of Deviance Table (Type II Wald chisquare tests)
##
## Response: FULLAUC
##                                Chisq Df Pr(>Chisq)
## log(PAUC + 1)                 2.6644  1  0.1026137
## PARINFTREAT                   5.5891  1  0.0180723 *
## SEX                           1.0547  1  0.3044333
## preSMI                        1.4134  1  0.2344942
## LENGTHRESID                   2.5699  1  0.1089134
## DOSE                          3.8052  1  0.0510929 .
## INFTYPE                       1.9123  1  0.1667064
## AGEATINF                     19.7368  1  8.887e-06 ***
## log(PAUC + 1):PARINFTREAT    11.8921  1  0.0005637 ***
## PARINFTREAT:AGEATINF          2.0272  1  0.1545011
## log(PAUC + 1):PARINFTREAT:SEX  0.6956  2  0.7062340
## ---
## Signif. codes:  0 '***' 0.001 '**' 0.01 '*' 0.05 '.' 0.1 ' ' 1
```

```
# Model fit
sim_residuals_glmmTMB <- simulateResiduals(m10, 1000)
plot(sim_residuals_glmmTMB)
```

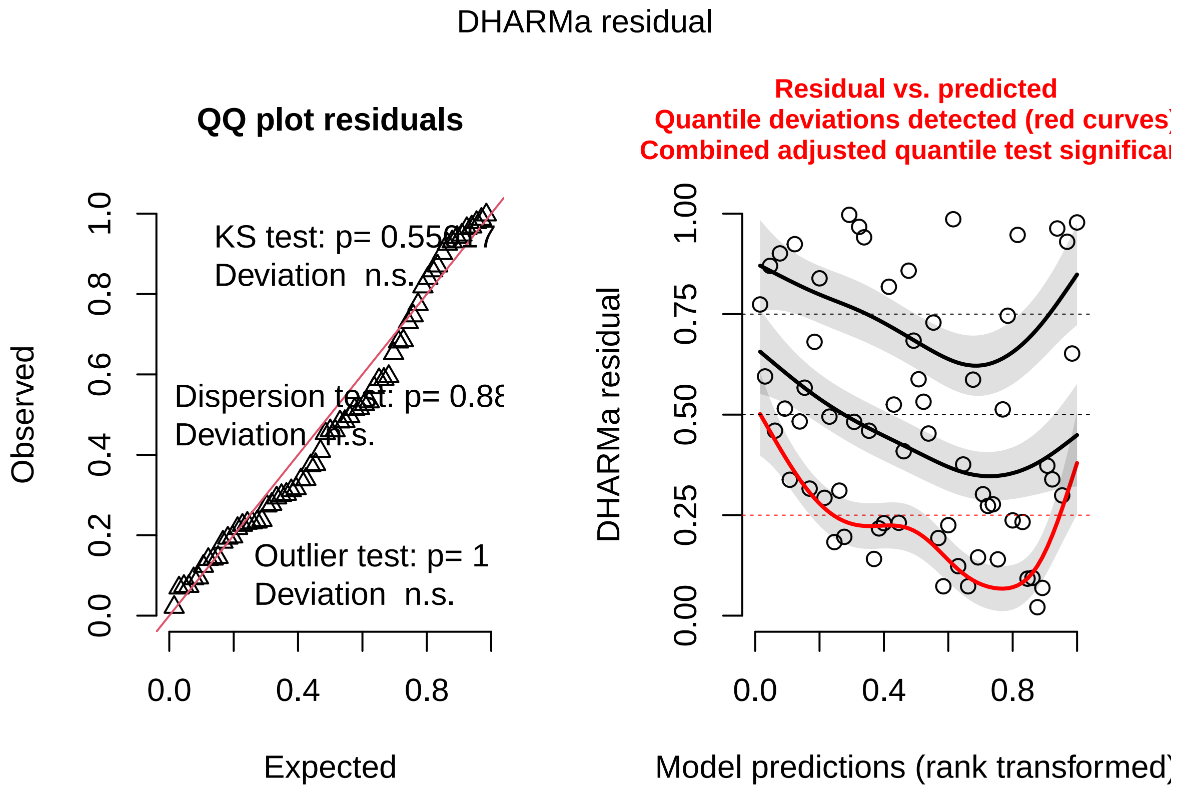

```
DHARMa::testDispersion(sim_residuals_glmmTMB)
```

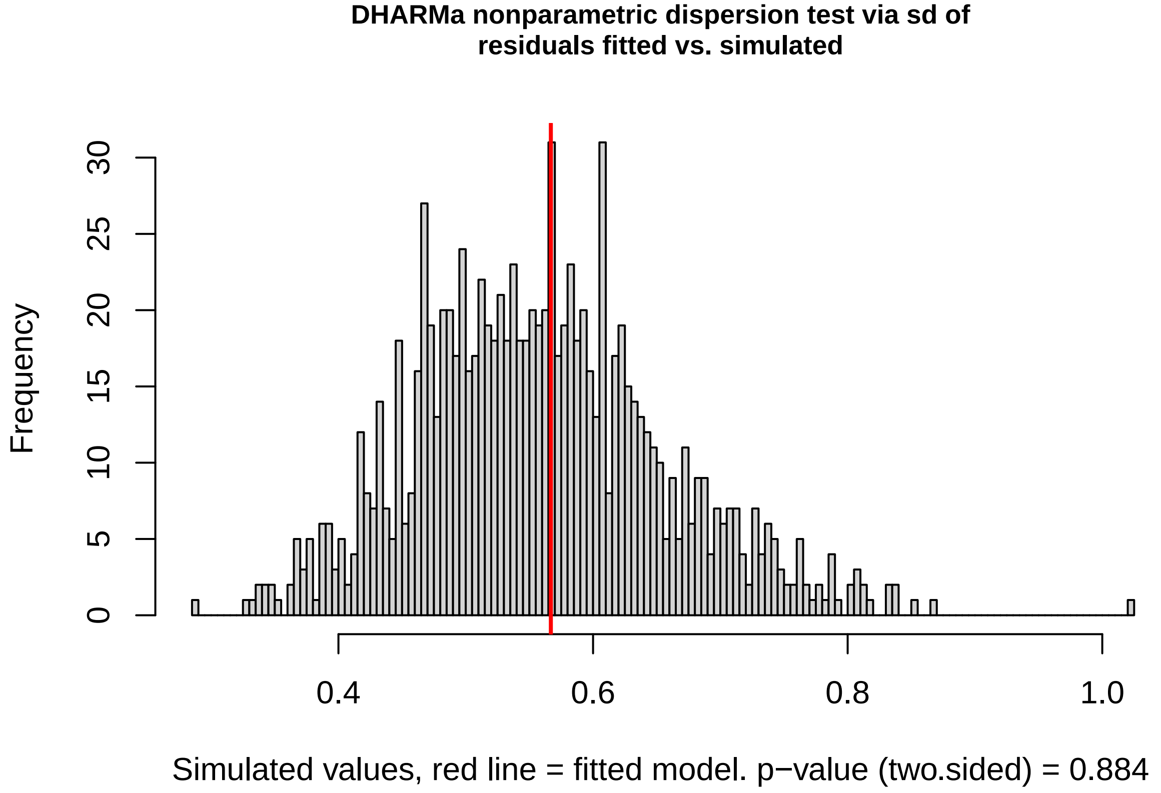

```
##
##  DHARMa nonparametric dispersion test via sd of residuals fitted vs.
```

```
##  simulated
##
## data:  simulationOutput
## dispersion = 1.0144, p-value = 0.884
## alternative hypothesis: two.sided
```

##### Post-hoc tests of the sex difference in the parent-offspring regression

These show that the results in m10 (Fig. 3 in the main text) are driven by the significant positive correlation between father AUC and offspring AUC, and no significant correlation exists between mother AUC and offspring AUC.

```r
# Full model
m_paternal <- glmmTMB(FULLAUC ~ log(PAUC + 1) + SEX + log(PAUC +
    1):SEX + DOSE + preSMI + LENGTHRESID + AGEATINF + (1 | MOTHERID),
    data = INDPARINFECTEDFATHER)

Anova(m_paternal)
```

```
## Analysis of Deviance Table (Type II Wald chisquare tests)
##
## Response: FULLAUC
##                     Chisq Df Pr(>Chisq)
## log(PAUC + 1)     13.0676  1  0.0003004 ***
## SEX                1.1217  1  0.2895527
## DOSE               0.2422  1  0.6226011
## preSMI             1.6009  1  0.2057741
## LENGTHRESID        2.3615  1  0.1243625
## AGEATINF          16.5660  1  4.699e-05 ***
## log(PAUC + 1):SEX  0.8030  1  0.3702068
## ---
## Signif. codes:  0 '***' 0.001 '**' 0.01 '*' 0.05 '.' 0.1 ' ' 1
```

```r
# Model fit
sim_residuals_glmmTMB <- simulateResiduals(m_paternal, 1000)
plot(sim_residuals_glmmTMB)
```

```
## qu = 0.25, log(sigma) = -3.108562 : outer Newton did not converge fully.
```

```
## qu = 0.5, log(sigma) = -2.516664 : outer Newton did not converge fully.
```

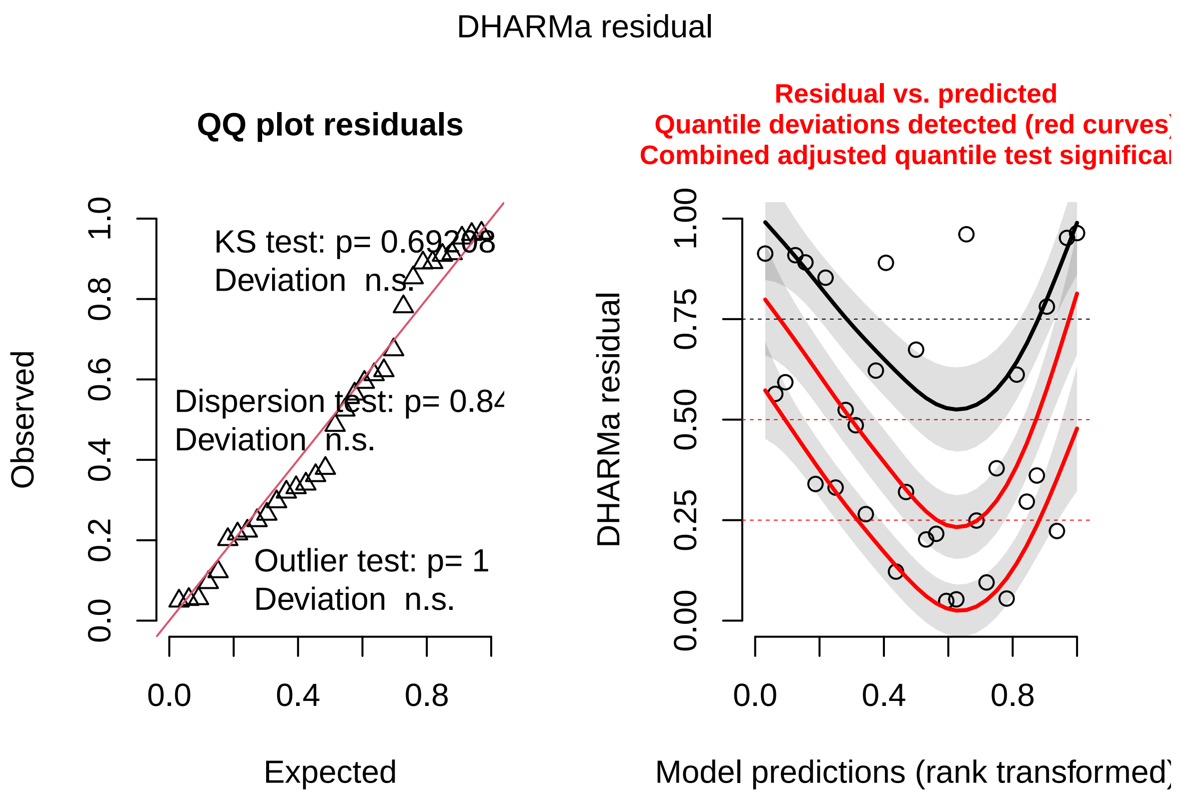

```
DHARMa::testDispersion(sim_residuals_glmmTMB)
```

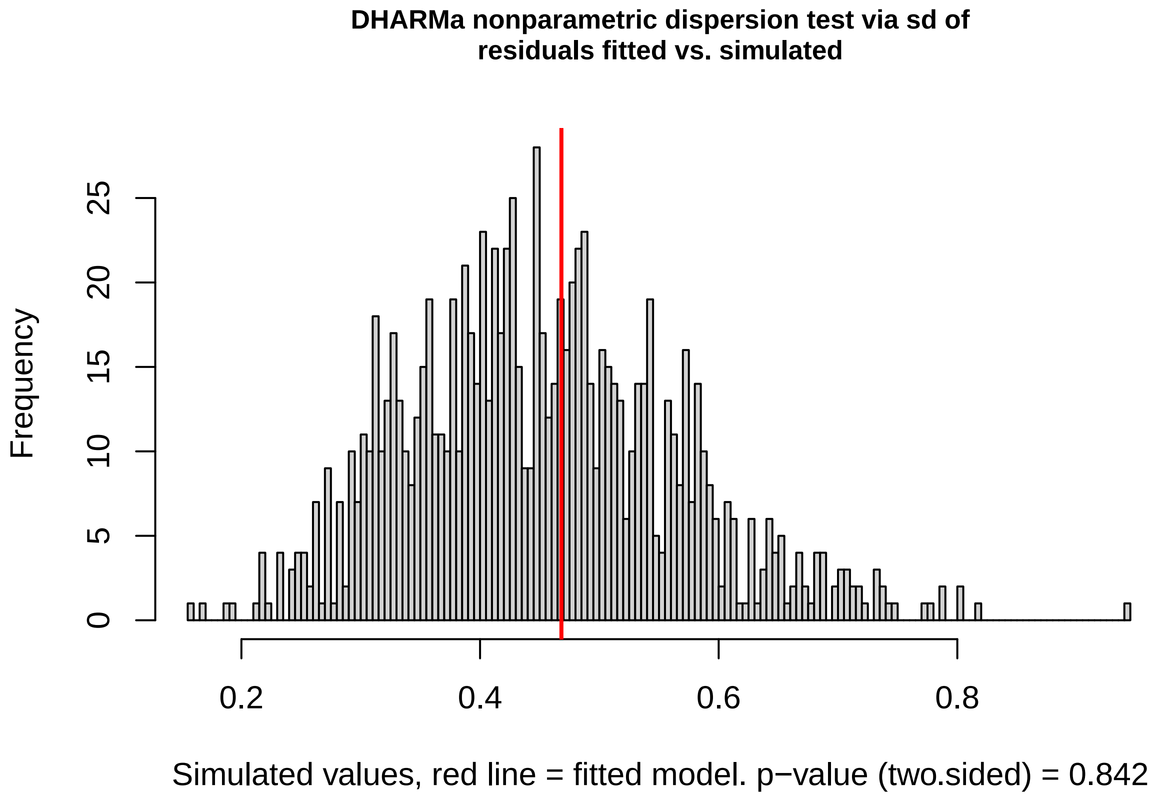

```
##
##  DHARMa nonparametric dispersion test via sd of residuals fitted vs.
```

```
##  simulated
##
## data:  simulationOutput
## dispersion = 1.038, p-value = 0.842
## alternative hypothesis: two.sided
```

```r
# Full model
m_maternal <- glmmTMB(FULLAUC ~ log(PAUC + 1) + SEX + DOSE +
    preSMI + LENGTHRESID + AGEATINF + log(PAUC + 1):SEX + (1 |
    MOTHERID), data = INDPARINFECTEDMOTHER)

Anova(m_maternal)
```

```
## Analysis of Deviance Table (Type II Wald chisquare tests)
##
## Response: FULLAUC
##                    Chisq Df Pr(>Chisq)
## log(PAUC + 1)     0.6303  1    0.42724
## SEX               0.4396  1    0.50732
## DOSE              1.6946  1    0.19299
## preSMI            0.3534  1    0.55220
## LENGTHRESID       0.4121  1    0.52092
## AGEATINF          4.7014  1    0.03014 *
## log(PAUC + 1):SEX 0.0063  1    0.93697
## ---
## Signif. codes:  0 '***' 0.001 '**' 0.01 '*' 0.05 '.' 0.1 ' ' 1
```

```r
# Model fit
sim_residuals_glmmTMB <- simulateResiduals(m_maternal, 1000)
plot(sim_residuals_glmmTMB)
```

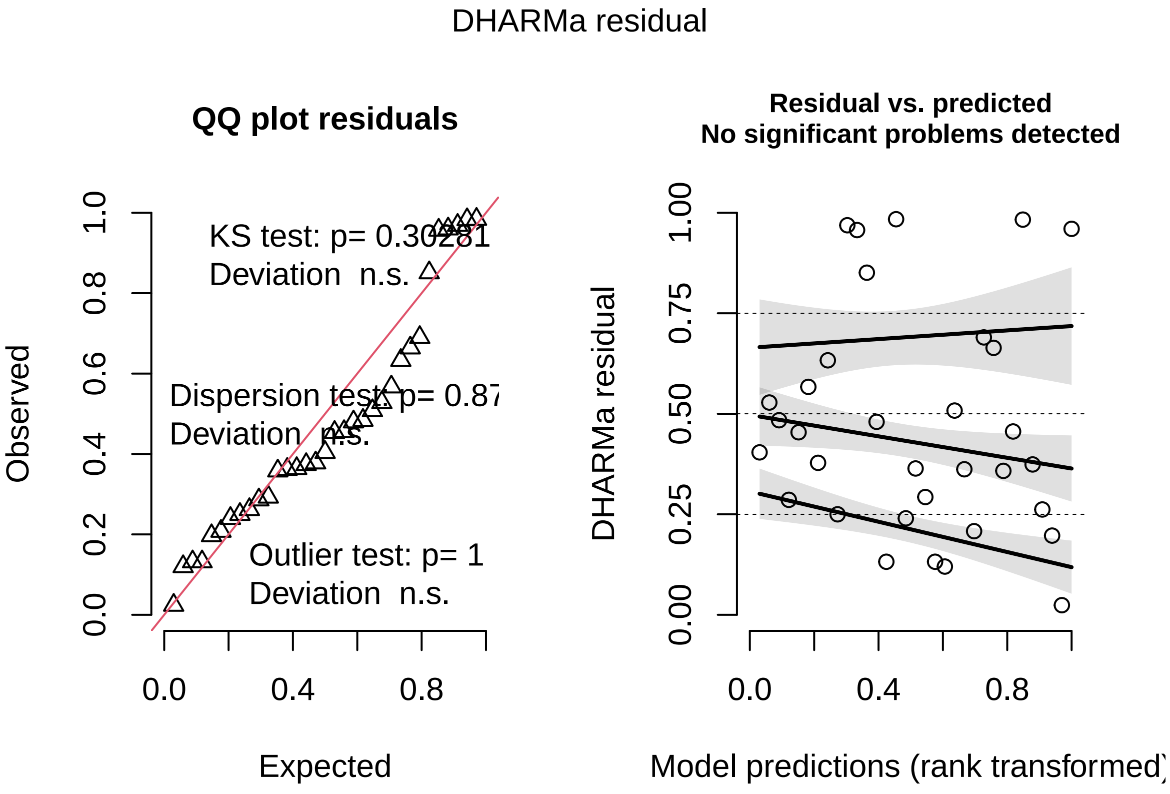

```r
DHARMa::testDispersion(sim_residuals_glmmTMB)
```

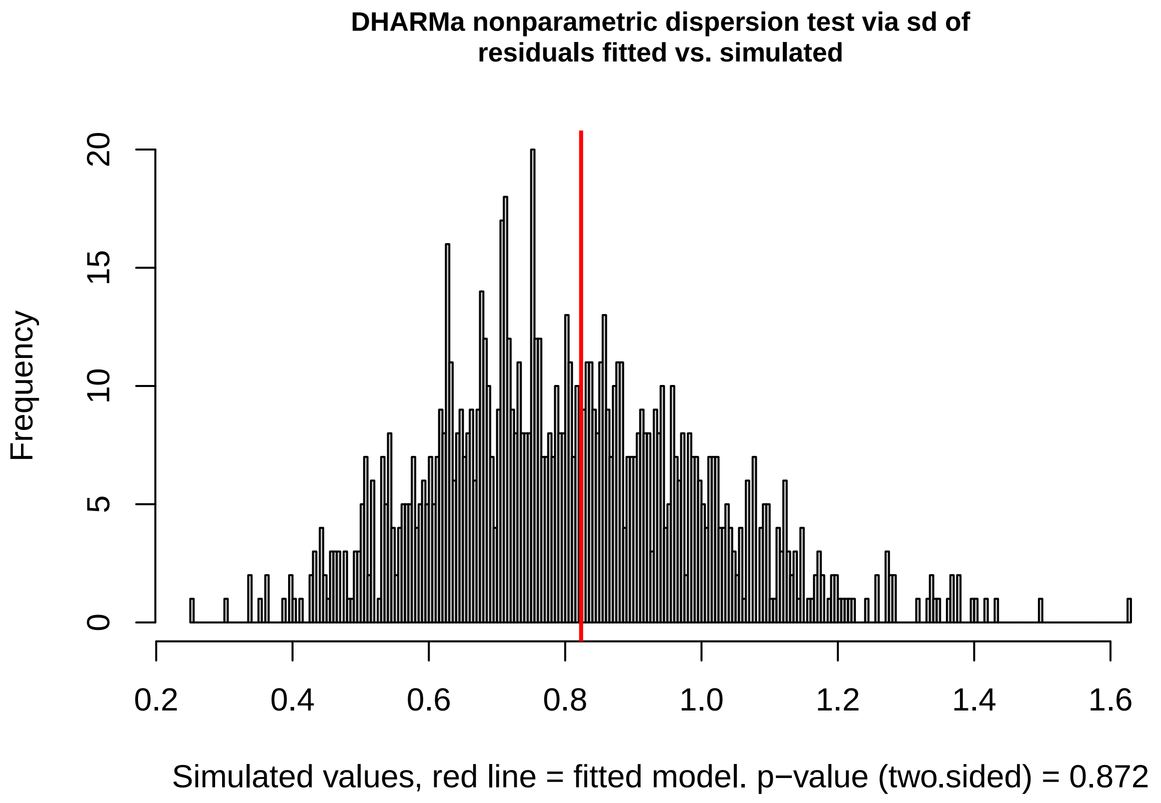

```
##
##  DHARMa nonparametric dispersion test via sd of residuals fitted vs.
##  simulated
##
## data:  simulationOutput
## dispersion = 1.0224, p-value = 0.872
## alternative hypothesis: two.sided
```

### Calculating the heritability of parasite resistance from parent-offspring regressions

#### Choosing the right representation of parent and offspring trait

Here we calculate the "half heritability" of parasite resistance from the regressions between parent infection integral and offspring infection integral. To compare between parents and offspring, the traits need to have the same distribution. We compared the distribution of the raw data, rescaled data, and log transformed data, and found that the rescaled data was the most similarly distributed across the two generations, as illustrated by the histograms. Below is code preparing the data for the analyses.

```r
# Subset INDPARINFECTED for infected fathers
INDPARINFECTEDFATHER <- subset(INDPARINFECTED, PARINFTREAT ==
    "FATHER")
# Subset INDPARINFECTED for infected mothers
INDPARINFECTEDMOTHER <- subset(INDPARINFECTED, PARINFTREAT ==
    "MOTHER")
```

```r
# Rescale parental AUC and recombined dataframe
INDPARINFECTEDMOTHER$PARAUC <- rescale(INDPARINFECTEDMOTHER$PAUC)
INDPARINFECTEDFATHER$PARAUC <- rescale(INDPARINFECTEDFATHER$PAUC)
INDPARINFECTED <- rbind(INDPARINFECTEDFATHER, INDPARINFECTEDMOTHER)

# create logged variables
INDPARINFECTED <- mutate(INDPARINFECTED, logFULLAUC = (log(FULLAUC +
    1)))
INDPARINFECTED <- mutate(INDPARINFECTED, logFAUC = (log(FAUC +
    1)))
INDPARINFECTED <- mutate(INDPARINFECTED, logMAUC = (log(MAUC +
    1)))
INDPARINFECTED <- mutate(INDPARINFECTED, logPAUC = (log(PAUC +
    1)))

# rescale so they are comparable
INDPARINFECTED$OFFAUC <- rescale(INDPARINFECTED$FULLAUC)
INDPARINFECTED$MUMAUC <- rescale(INDPARINFECTED$MAUC)
INDPARINFECTED$DADAUC <- rescale(INDPARINFECTED$FAUC)
INDPARINFECTED$REAGE <- rescale(INDPARINFECTED$AGEATINF)

# remove one outlier
IND2 <- subset(INDPARINFECTED, FULLAUC < 1000)
```

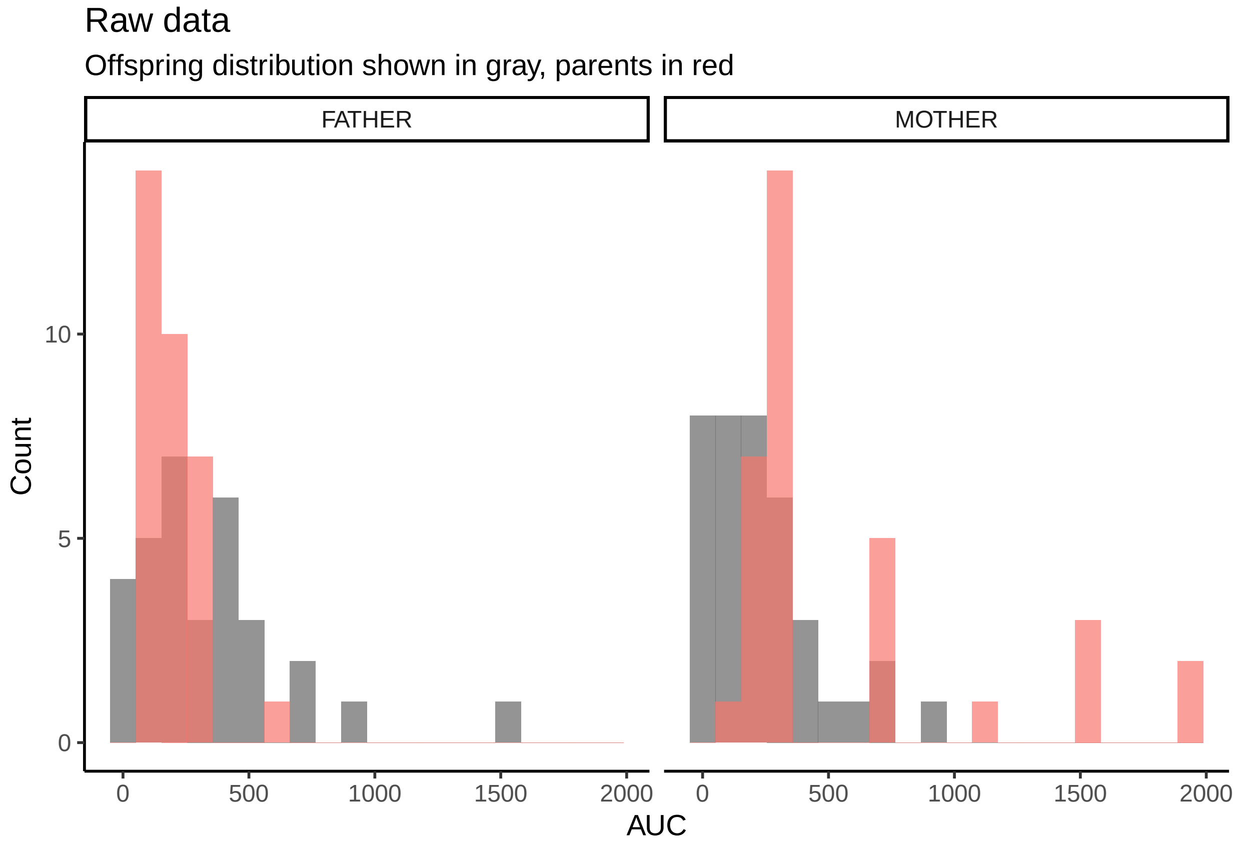

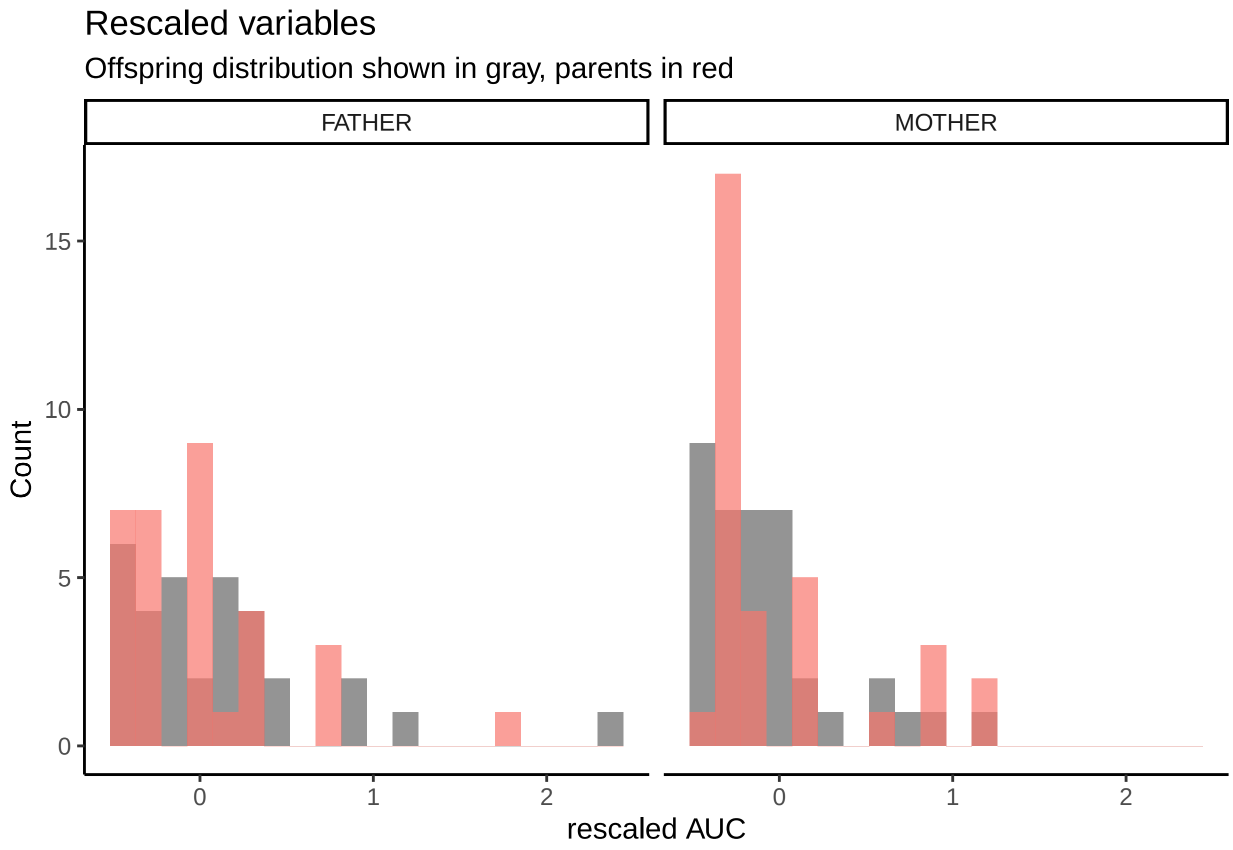

```
## Warning: Removed 5 rows containing non-finite outside the scale range
## (`stat_bin()`).
```

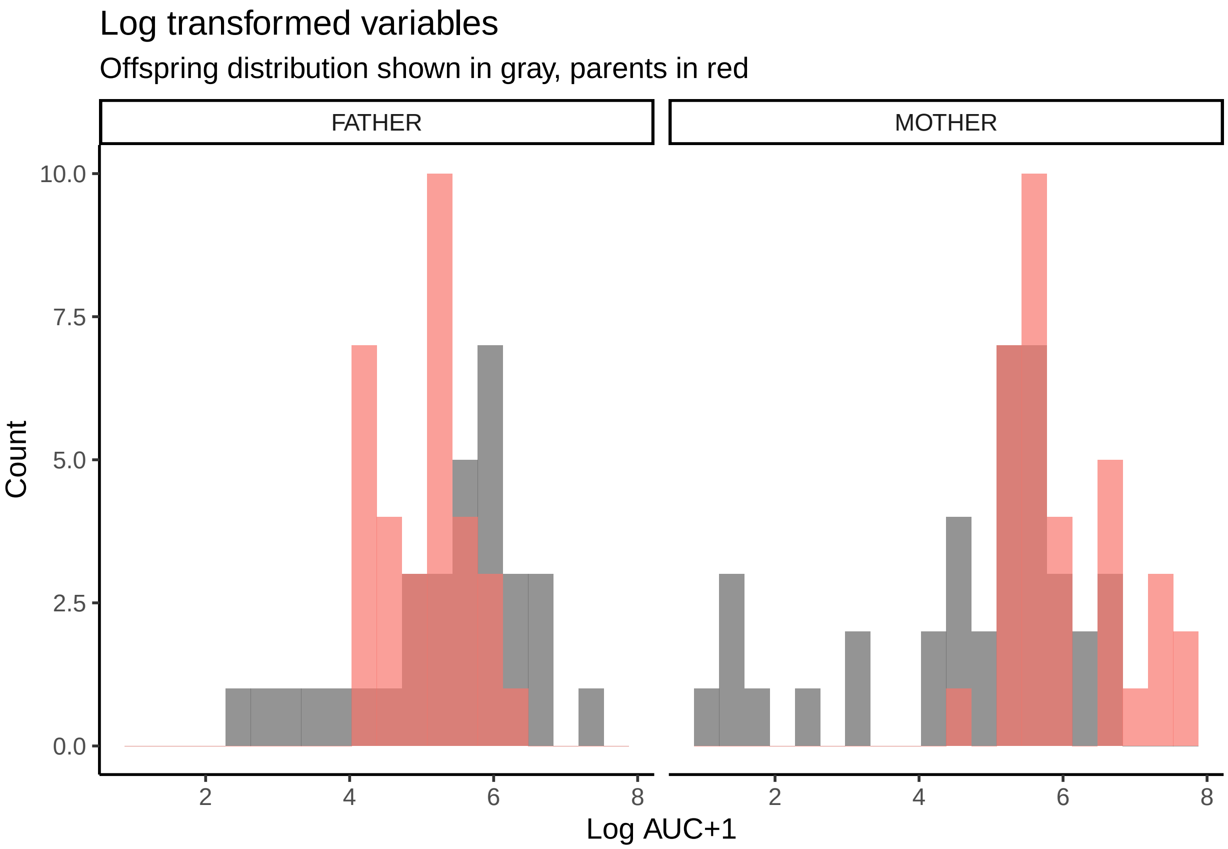

#### Obtaining slope - i.e., heritability - estimates

We used offspring infection integral (rescaled) as the response variable. As explanatory variables, we used offspring age at infection (rescaled; 'REAGE') because previous results indicated it explained a highly significant portion of the variation in offspring infection integral. We included mother ID as a random effect to control for the fact that several pairs contributed more than one offspring. Because there is one conspicuous outlier we repeated the analysis without it to check its effects.

##### Father-offspring regression

The father-offspring regression slope is 0.69±0.13, and is highly significant. After removal of the outlier, the estimate is 0.53±18, and remains highly significant.

```
# Rescaled variables for Father:
dadh <- glmmTMB(OFFAUC ~ DADAUC + REAGE + (1 | MOTHERID), data = INDPARINFECTED)
summary(dadh)$coefficients$cond
```

```
##               Estimate Std. Error  z value     Pr(>|z|)
## (Intercept) 0.08551053 0.06284783 1.360596 1.736413e-01
## DADAUC      0.64543436 0.12858085 5.019677 5.175843e-07
## REAGE       0.52173965 0.12412777 4.203247 2.631133e-05
```

```
father_halfh <- visreg(dadh, "DADAUC", gg = T) + stat_regline_equation(label.y = 1.5,
    aes(label = after_stat(eq.label))) + scale_y_continuous("Offspring infection integral") +
    scale_x_continuous("Father infection integral") + theme_classic()
print(father_halfh)
```

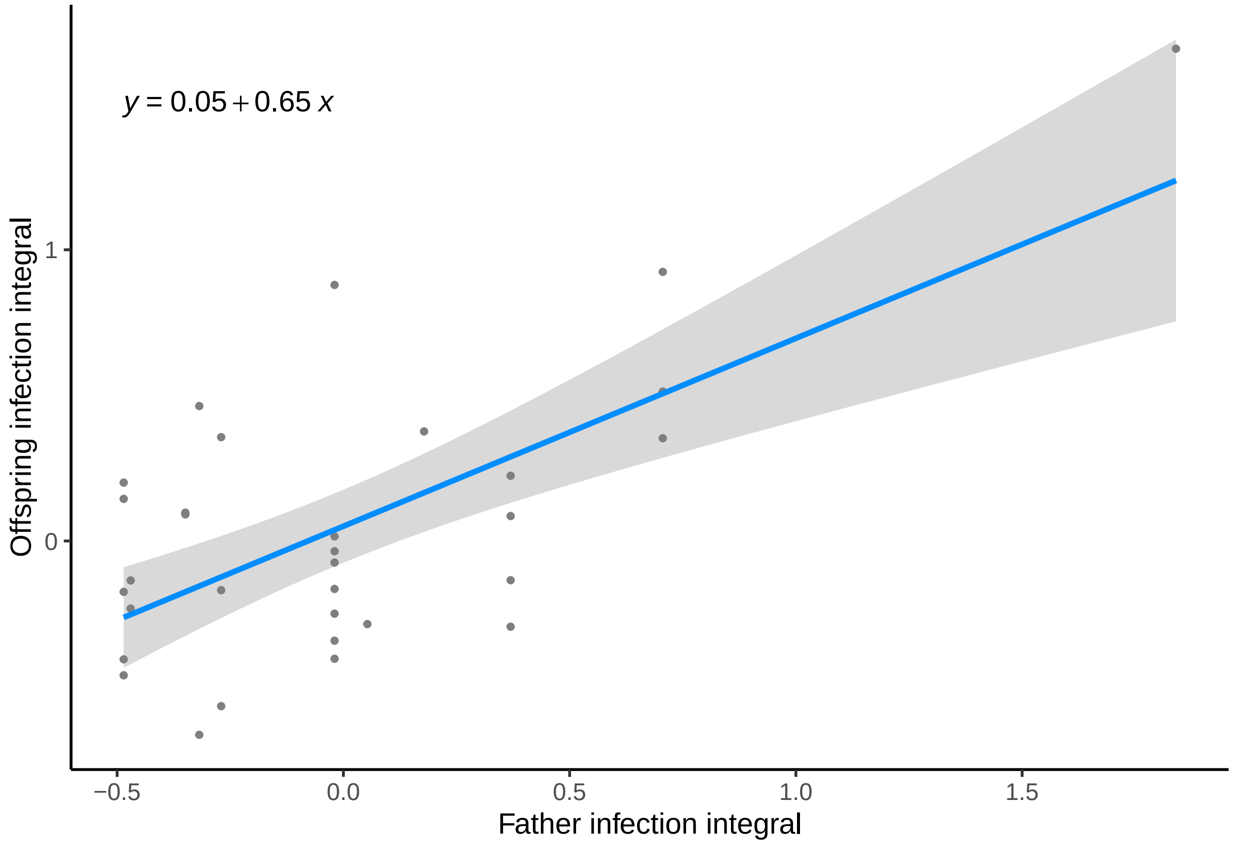

```r
# Without the outlier:
dadho <- glmmTMB(OFFAUC ~ DADAUC + REAGE + (1 | MOTHERID), data = IND2)
summary(dadho)$coefficients$cond
```

```
##                Estimate Std. Error  z value    Pr(>|z|)
## (Intercept) 0.05810312 0.06172088 0.941385 0.346507574
## DADAUC      0.42221975 0.16679185 2.531417 0.011360267
## REAGE       0.42301233 0.12881811 3.283796 0.001024192
```

```r
father_halfho <- visreg(dadho, "DADAUC", gg = T) + stat_regline_equation(label.y = 0.5,
    aes(label = after_stat(eq.label))) + scale_y_continuous("Offspring infection integral (rescaled)") +
    scale_x_continuous("Father infection integral (rescaled)") +
    theme_classic()
print(father_halfho)
```

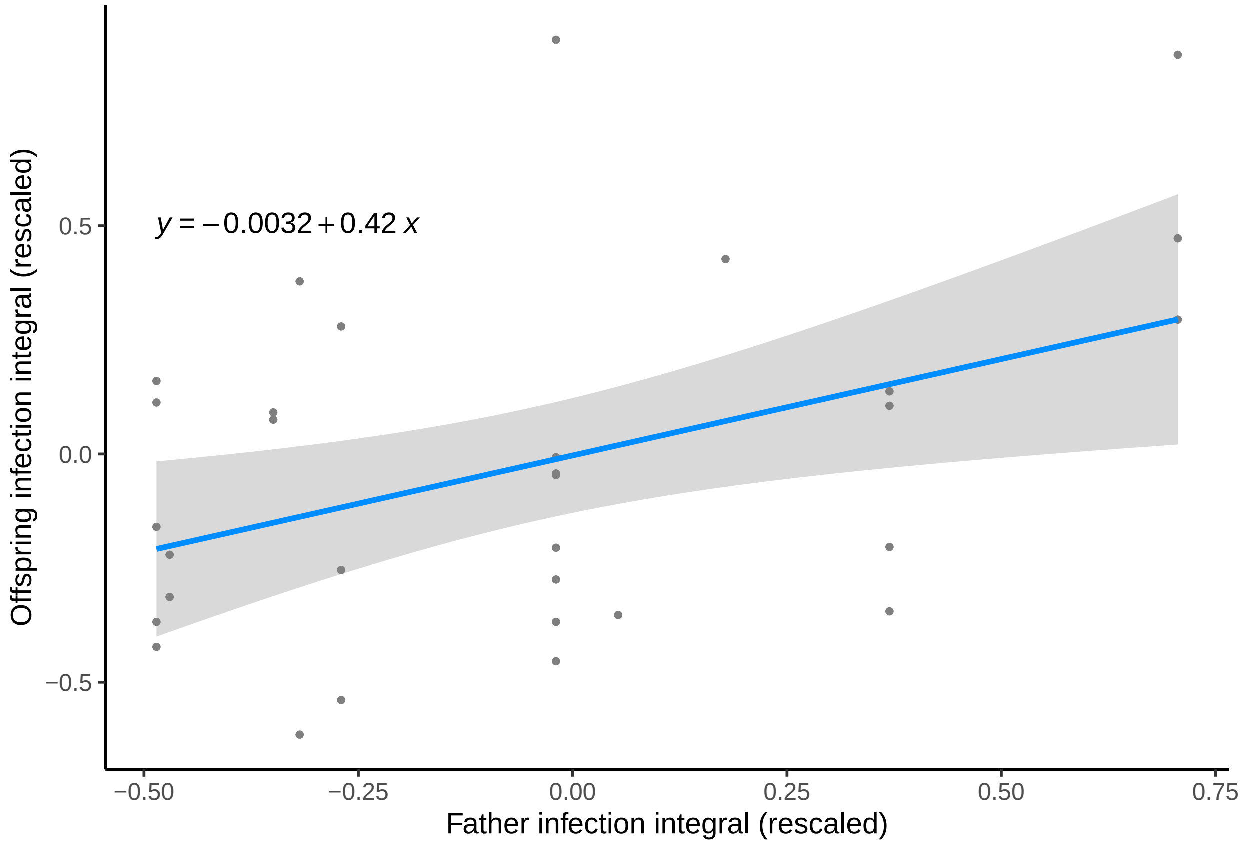

###### Mother-offspring regression

There is no outlier in this dataset. The estimate is -0.13±0.17, and the slope is not significantly different from 0.

```
# Rescaled variables for Mother:
mumh <- glmmTMB(OFFAUC ~ MUMAUC + REAGE + (1 | MOTHERID), data = INDPARINFECTED)
summary(mumh)$coefficients$cond
```

```
##                Estimate Std. Error    z value   Pr(>|z|)
## (Intercept) -0.05650763 0.06958131 -0.8121093 0.41672893
## MUMAUC      -0.16126123 0.14193528 -1.1361603 0.25588949
## REAGE        0.25568276 0.13520260  1.8911084 0.05860988
```

```
mother_halfh <- visreg(mumh, "MUMAUC", gg = T) + stat_regline_equation(label.y = 0.5,
    aes(label = after_stat(eq.label))) + scale_x_continuous("Mother infection integral (rescaled)") +
    scale_y_continuous("Offspring infection integral (rescaled)") +
    theme_classic()
print(mother_halfh)
```

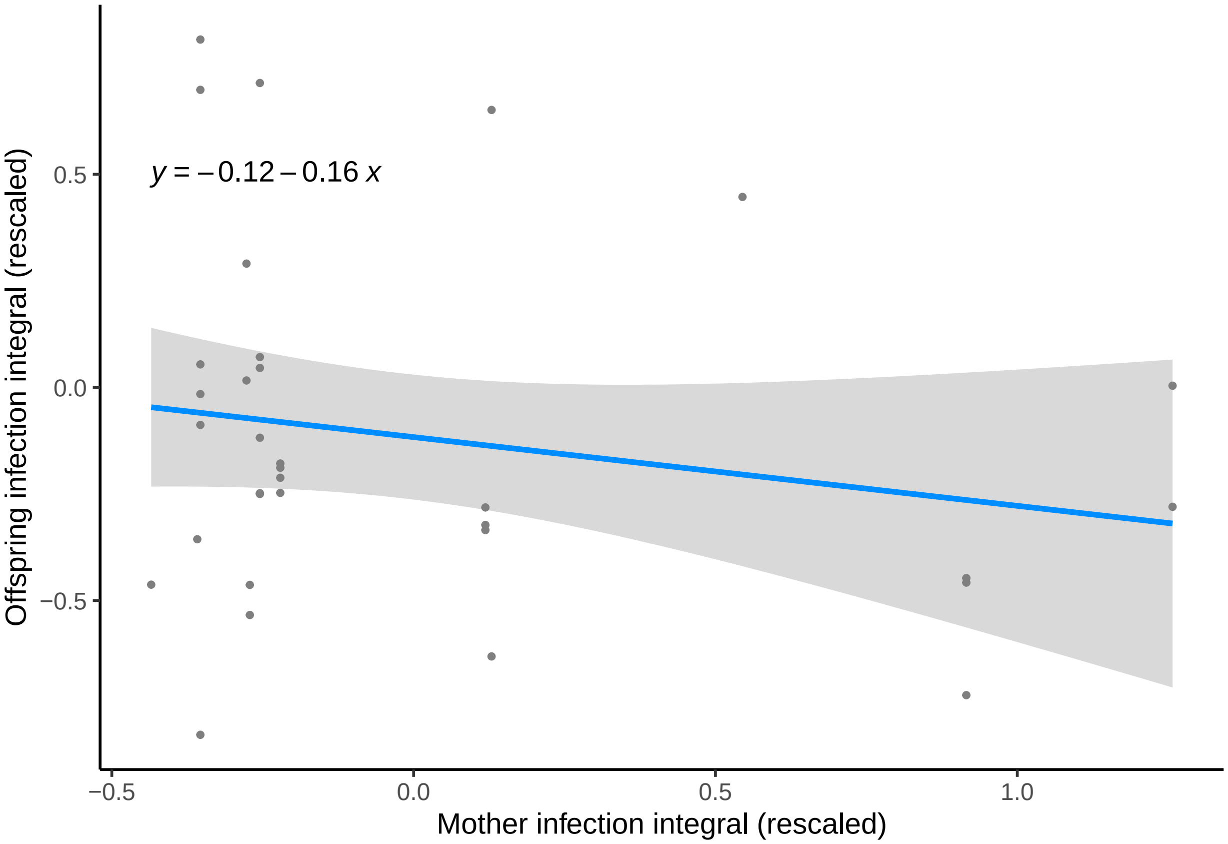

$y = -0.12 - 0.16\,x$
Offspring infection integral (rescaled)
0.5
0.0
−0.5
−0.5
0.0
0.5
1.0
Mother infection integral (rescaled)
